# Immature *C. elegans* motor neurons control early embryo behavior via both synaptic and non-synaptic GABA release

**DOI:** 10.1101/2025.11.19.689106

**Authors:** James Marvel-Coen, Evan Ardiel, Jian Zhao, Stephen Nurrish, Joshua M. Kaplan

## Abstract

Pre-natal brain activity has long lasting effects on subsequent neurodevelopment. It is unclear if early brain activity is dominated by cell intrinsic, synaptic, or non-synaptic mechanisms. We address this question by analyzing *C. elegans* embryo behavior in *snf-11* mutants, which lack a plasma membrane GABA re-uptake pump (orthologous to GAT1). At 510-570 minutes post-fertilization, embryo motion was transiently and potently inhibited in *snf-11* GAT1 mutants, which precedes formation of most nerve ring synapses. This transient motion inhibition requires GABA synthesis in DD motor neurons and UNC-49 GABA_A_ receptors in body muscles. When motion inhibition occurs, DD neurons have not yet completed neurite outgrowth. Genetic analysis suggests that motion inhibition was mediated by both synaptic and tonic GABA release from DD motor neurons. These results suggest that DD neurons control embryo behavior prior to completing their developmental maturation.

**Significance Statement:** Little is known about how prenatal circuits control embryo behavior. We show that the motion of early *C. elegans* embryos is transiently inhibited by immature GABAergic motor neurons that have not yet completed neurite outgrowth. GABA’s inhibitory effect on early embryo behavior occurs before most synapses have formed in the worm’s central neuropil. For this early behavior, we find that synaptic GABA release plays a minor role while non-synaptic mechanisms predominate. Our results suggest that non-synaptic forms of GABA transmission play a significant role in prenatal circuits.

## Introduction

Many results suggest that early events in brain development play a pivotal role in post-natal developmental outcomes. Development of some circuits is restricted to particular times early in development (i.e., critical periods) (1–3). Maternal infection, maternal drug exposure, and prematurity are all associated with increased risk for neurodevelopmental disorders (4–6). These and other results suggest that early brain circuits may be particularly vulnerable to genetic and environmental perturbation. Although behavior and circuit function have been intensively investigated, the vast majority of these studies have used post-natal samples, presumably due to the inaccessibility of pre-natal tissues. Consequently, very little is known about prenatal behaviors and the circuit mechanisms producing them. Several important questions about early brain development remain to be answered. What are the first behaviors exhibited by embryos? When do neurons first assume control of embryo behavior? Do neurons control behavior prior to completing their developmental maturation? Do immature neurons encode behavior by mechanisms distinct from those used in post-natal circuits?

In many systems, embryos begin exhibiting behaviors before birth. *C. elegans* embryos begin moving shortly before the 2-fold stage; *Drosophila* embryos at stage 16, 800-900 mpf; chick embryos at stage 21 (∼3.5 days), and human embryos at 8 weeks (7–9). Embryonic movement is necessary for circuit formation and even survival in invertebrate model systems including *Drosophila* and *C. elegans* (8, 10–12).

Although prenatal behavior is widely observed, when synapses form and assume control of embryo behavior has not been determined in most systems. *In vitro*, developing neurons can release neurotransmitters from growth cones (13, 14). In mice, motor axons reach muscles by E12.5 but clustered acetylcholine receptors (AChR) are not observed until E14.5 (15). *In vivo* recordings of pyramidal neurons in fetal mice indicate synaptic activity commences by E14.5 (16). In contrast, perinatal rat (day E18-P1) hippocampal slices exhibited both evoked and tonic GABA activated currents, which were not blocked by botulinum toxin nor by a general voltage-gated calcium channel antagonist (CdCl_2_), suggesting that these were mediated by non-synaptic mechanisms (17).

The timing of embryonic neurodevelopment has also been extensively analyzed in *C. elegans* (Fig. 1A)*. C. elegans* embryos develop inside an eggshell before hatching as a first stage larva, at ∼800 minutes post-fertilization (mpf). Neurodevelopmental landmarks occur at stereotyped times of *C. elegans* embryonic development. Neuron births occur in two waves (230-290 mpf and 400 mpf) in the cell lineage (7). Axon outgrowth in the nerve ring (the major neuropil in the worm’s head) is completed at 460 mpf (18). Several pre-synaptic proteins are localized to the nerve ring at ∼400 mpf (19). At 550 mpf, small neuromuscular junctions (NMJs) are observed in electron micrographs of the dorsal nerve cord whereas ventral NMJs are not yet seen at this time (20).

**Figure 1.**
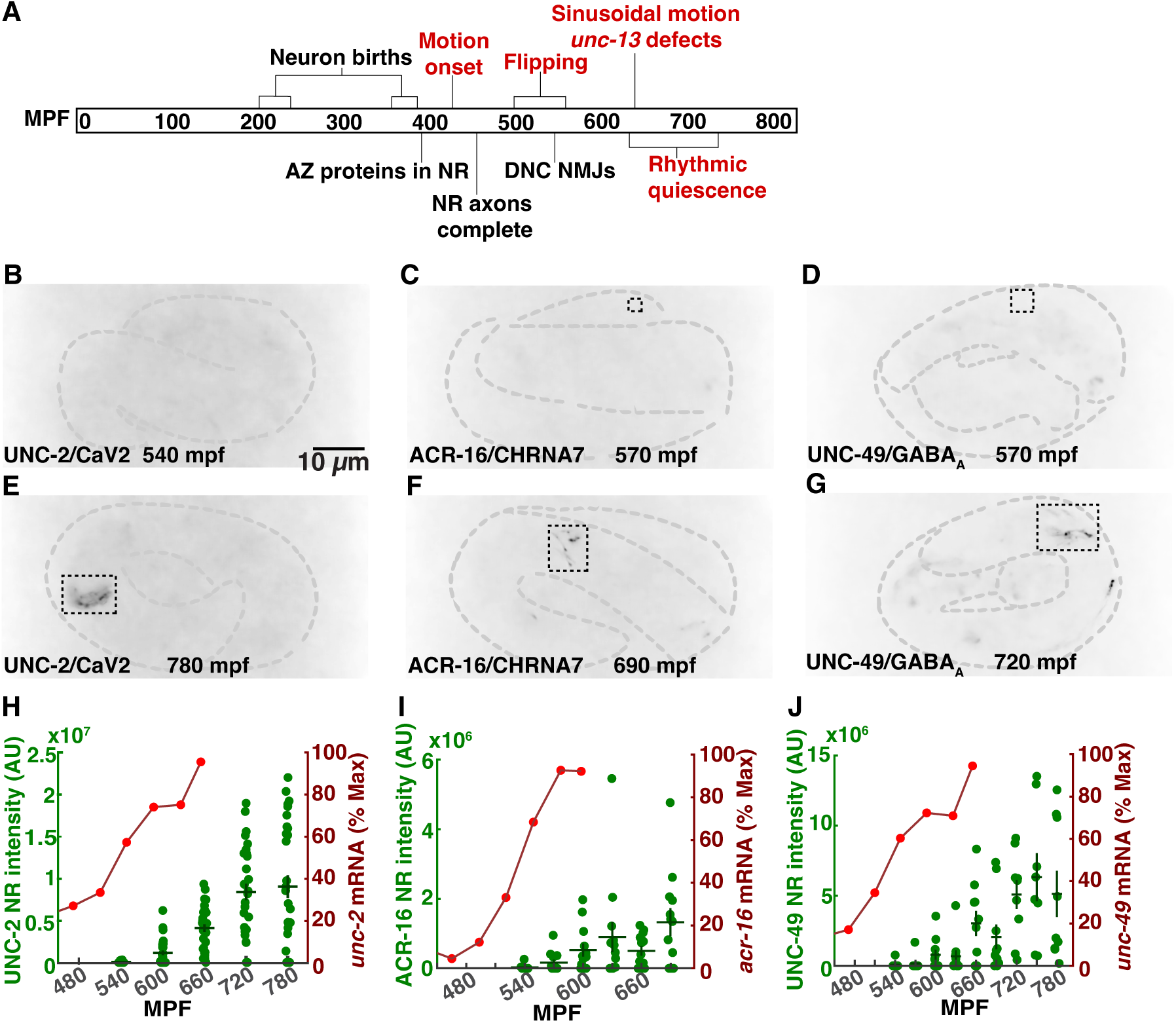
Synaptic ion channels localize to the nerve ring after 600 mpf. A) An overview of *C. elegans* embryonic neurodevelopment. Anatomical (black) and behavioral (red) milestones are indicated. B-G) Representative images of early (B-D) and late (E-G) embryos expressing UNC-2/Ca_V_2(nu569 mNG) (A,D), ACR-16/CHRNA7(nu718 mNG), and UNC-49/GABA_A_R(nu829 mNG) are shown. In each case, nerve ring signal (within the dashed rectangles) is observed in the late (E-G) but not early (B-D) embryos. Developmental time point is indicated for each image. UNC-2/CaV2 is expressed in all neurons (E); ACR-16/CHRNA7 nerve ring signal (F) is primarily in AVA neurons; and UNC-49/GABA_A_R nerve ring signal (G) occurs in four regions (two of which are shown in this image), which likely represents clustered body muscle receptors at RME neuron NMJs. (H-J) Summary data for mean nerve ring integrated fluorescence intensity and peak normalized mRNA levels (Boeck et al., 2016) versus developmental time are shown for UNC-2/CaV2 (H), ACR-16/CHRNA7 (I), and UNC-49/GABA_A_R (J). The RNA sequencing data were taken from a prior publication (37, 76). Error bars indicate SEM.

In parallel to this developmental timeline, *C. elegans* embryos also exhibit a stereotyped pattern of behavioral maturation (Fig. 1A). Spontaneous apparently uncoordinated movement (termed twitches) appear at 430 mpf (7, 12). At 530-570 mpf, embryos exhibit an alternating pattern of all ventral and all dorsal body bends, hereafter designated flipping behavior (21). Flipping is not significantly disrupted in *unc-13* mutants (21), which nearly completely lack SV exocytosis (22), suggesting that flipping is not mediated by a synaptic circuit. Finally, at 650-700 mpf, a mature pattern of sinusoidal motion and rhythmic quiescent bouts emerge, both of which are profoundly inhibited in *unc-13* mutants (indicating that they are driven by synaptic circuits) (21).

To further address the mechanisms controlling early brain activity, we analyzed the impact of the neurotransmitter γ-aminobutyric acid (GABA) on *C. elegans* embryos. For several reasons, GABA is a particularly interesting case to consider. In mice, GABA plays a crucial role in early circuit formation (23–25). Inhibitory signaling from GABAergic neurons is required for the onset of plasticity during the critical period in mouse visual cortex development, and mutations in MeCP2, associated with Rett syndrome, alter visual cortex development by increasing GABA transmission (2, 11, 26). The GAT1 GABA re-uptake pump and GABRB3 GABA_A_ receptors are risk genes for autism spectrum disorder (ASD) (27–29), which is thought to arise from changes in fetal brain development (30–32). For these reasons, there is significant interest in understanding how and when GABA release occurs in early brain development.

Here we further investigate when synaptic circuits first form in *C. elegans* embryos. We find that many nerve ring synapses form after 600 mpf, and that an earlier embryo behavior (flipping) is controlled by both synaptic and non-synaptic GABA transmission. These results suggest non-synaptic GABA release plays an important role in early brain activity.

## Results

### Synaptic ion channels localize to the nerve ring at approximately 600 mpf

Prior studies provide conflicting conclusions about when functional synaptic circuits first form in *C. elegans* embryos. Depending on how synapses were assayed, estimates for when synapses form ranged from 400 mpf (19), to 550 mpf (20), to 650 mpf (21).

To further address this question, we asked when several synaptic ion channels are localized to the nerve ring (a large circumferential axon bundle in the worm’s head that contains many synaptic connections). Trafficking of large multimeric ion channels (e.g. ionotropic glutamate receptors) to the cell surface takes many hours due to their slow protein folding and subunit assembly in the endoplasmic reticulum (33, 34). Consequently, we reasoned that ion channels likely represent the last components to be localized to synapses. We analyzed the localization of endogenously expressed UNC-2 Ca_V_2 (a voltage-activated calcium channel), UNC-49 GABA_A_R (a GABA activated chloride channel), and ACR-16 CHRNA7 (a nicotinic acetylcholine receptor) in the nerve ring of 540 to 660 mpf embryos, using mNeonGreen (mNG)-tagged CRISPR alleles of each gene (Fig. 1B-J). Electrophysiological recordings show that the mNG tags did not impair the function of UNC-2 (35), UNC-49, and ACR-16 channels (*SI Appendix,* Fig. S1 A-D). All three channels arrived in the nerve ring at a similar time, starting at ∼600 mpf (Fig. 1B-J), which was ∼1-3 hours after mRNA expression onset (defined as the time when expression reached 20% of the peak value) (Fig. 1H-J) (36–38). By contrast, mNG tagged NRX-1 and NLG-1 (pre- and post-synaptic adhesion molecules respectively) arrived in the nerve ring significantly earlier than these ion channels (*SI Appendix,* Fig. S2), consistent with the earlier arrival times reported for other synaptic proteins (19). These results imply that some essential synaptic proteins do not appear in the nerve ring until 600 mpf or later, which coincides with when motion defects are first observed in *unc-13* mutants (21).

### GABA signaling decreases early embryo motion

To assess GABA function in early embryos, we analyzed the behavior of *snf-11* mutants. SNF-11 is a plasma membrane GABA reuptake pump (orthologous to GAT1) (39, 40). Thus, *snf-11* GAT1 mutants are predicted to have exaggerated GABA signaling. Embryo motion was assessed by frame subtraction of bright field images (21). Using this assay, we found that *snf-11* mutant embryos exhibit a unique behavioral defect whereby motion is dramatically decreased at a consistent developmental time point (∼510-570 mpf) before abruptly returning to a wild type pattern thereafter (Fig. 2A). Motion inhibition was quantified by measuring the maximal decrease in motion rate in individual 480-620 mpf embryos (as detailed in the Experimental Methods). This phenotype was observed in mutants containing three independent *snf-11* GAT1 loss of function alleles (Fig. 2B). This *snf-11* GAT1 mutant motion defect was rescued by transgenes expressing SNF-11 in either muscles or neurons (Fig. 2C), both of which express the endogenous *snf-11* gene (36). These results confirm that decreased embryo motion is a consequence of *snf-11* GAT1 inactivation. Because *snf-11* encodes a GABA re-uptake pump, we next asked if GABA synthesis is required for the reduced motion of *snf-11* embryos. Consistent with this idea, a mutation inactivating *unc-25*, which encodes the GABA biosynthetic enzyme glutamic acid decarboxylase (GAD), restores wild type motion in *snf-11* embryos (Fig. 2F). By contrast, *unc-25* GAD single mutants exhibited a significant increase in motion at ∼530-570 mpf relative to wild type (Fig. 2D and E). Collectively, these results suggest that decreased motion of *snf-11* GAT1 mutant embryos results from exaggerated GABA signaling.

**Figure 2.**
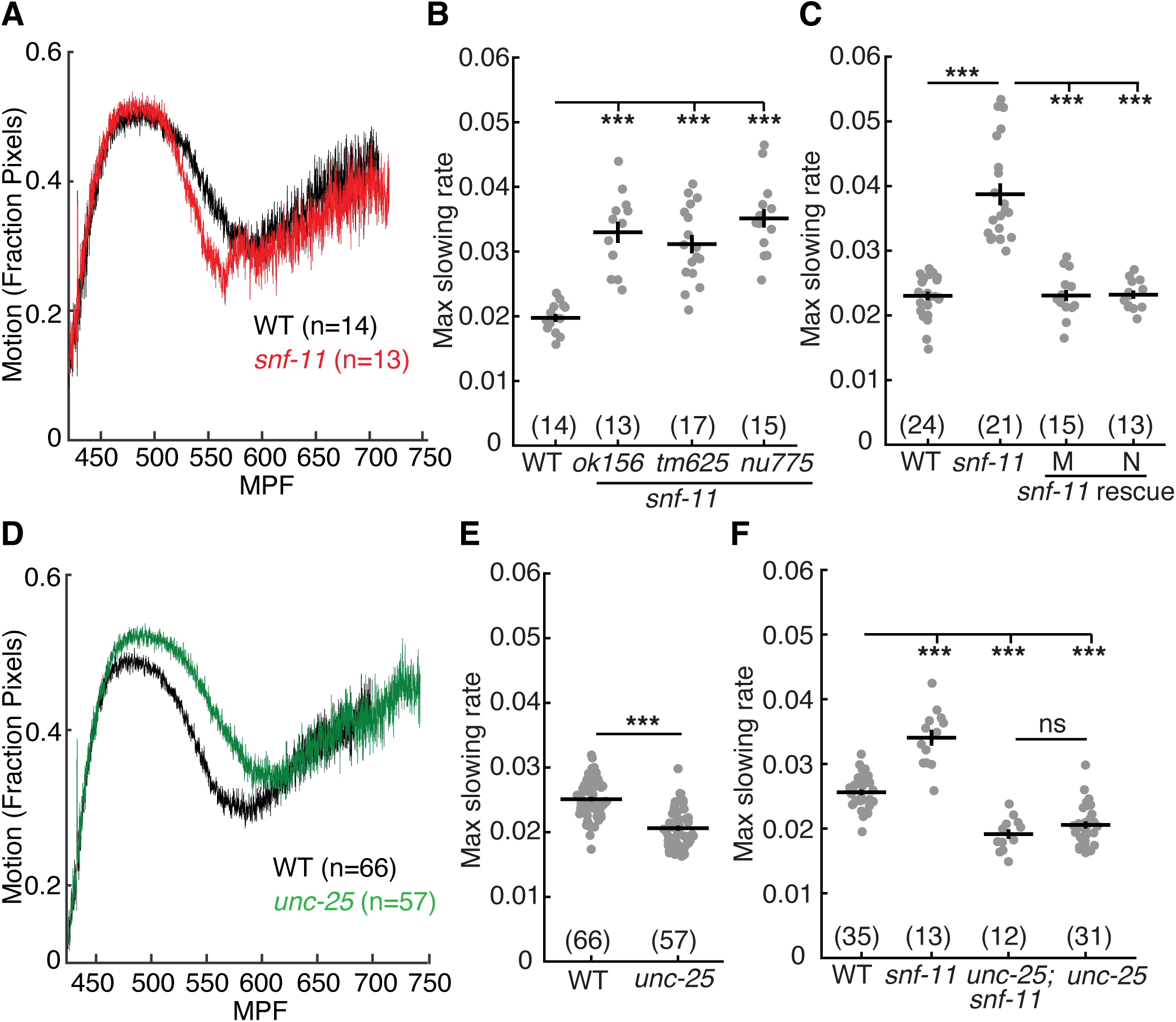
GABA transiently inhibits embryo motion. Average motion traces (A,D) and maximal slowing rate of 480-620 mpf embryos (B,C,E,F) are shown for the indicated genotypes. (A) Embryo motion was transiently inhibited in *snf-11(ok156)* GAT1 embryos (red) compared to WT controls (black). B) Three independent *snf-11* GAT1 loss of function alleles caused similar decreases in motion rate. C) The decreased motion defect in *snf-11(ok156)* mutants was rescued by transgenes restoring SNF-11 expression in muscles (M) or in neurons (N). D-E) Increased embryo motion was observed in *unc-25(nu836)* GAD (green) embryos compared to WT controls (black). F) The *snf-11* GAT1 decreased motion defect was eliminated in *unc-25; snf-11* double mutants. Sample sizes for each genotype are indicated in all figure panels. Values that differ significantly are indicated (ns, not significant; ***, *p* <0.001). Error bars indicate SEM.

### GABA inhibits embryo flipping behavior

The brief inhibition of motion observed in *snf-11* GAT1 mutants occurs when behavior is dominated by embryo flipping (21). To confirm that *snf-11* GAT1 mutations disrupt flipping, we used lightsheet microscopy to analyze *snf-11* GAT1 embryo postures, as described (21). Embryo postures were calculated by tracking the location of paired hypodermal skin cell that run along the left and right sides of the body. As previously described, ∼570 mpf wild type embryos regularly flip between full dorsal and full ventral coils (Fig. 3A) (21). By contrast, *snf-11* mutant embryos (at 570 mpf) adopted a single prolonged coiled posture (Fig. 3B). Taken together, these results suggest that exaggerated GABA signaling in *snf-11* GAT1 mutants dramatically (albeit transiently) suppresses early embryo flipping behavior. Hereafter, we refer to the *snf-11* mutant motion defects as inhibition of embryo flipping.

**Figure 3.**
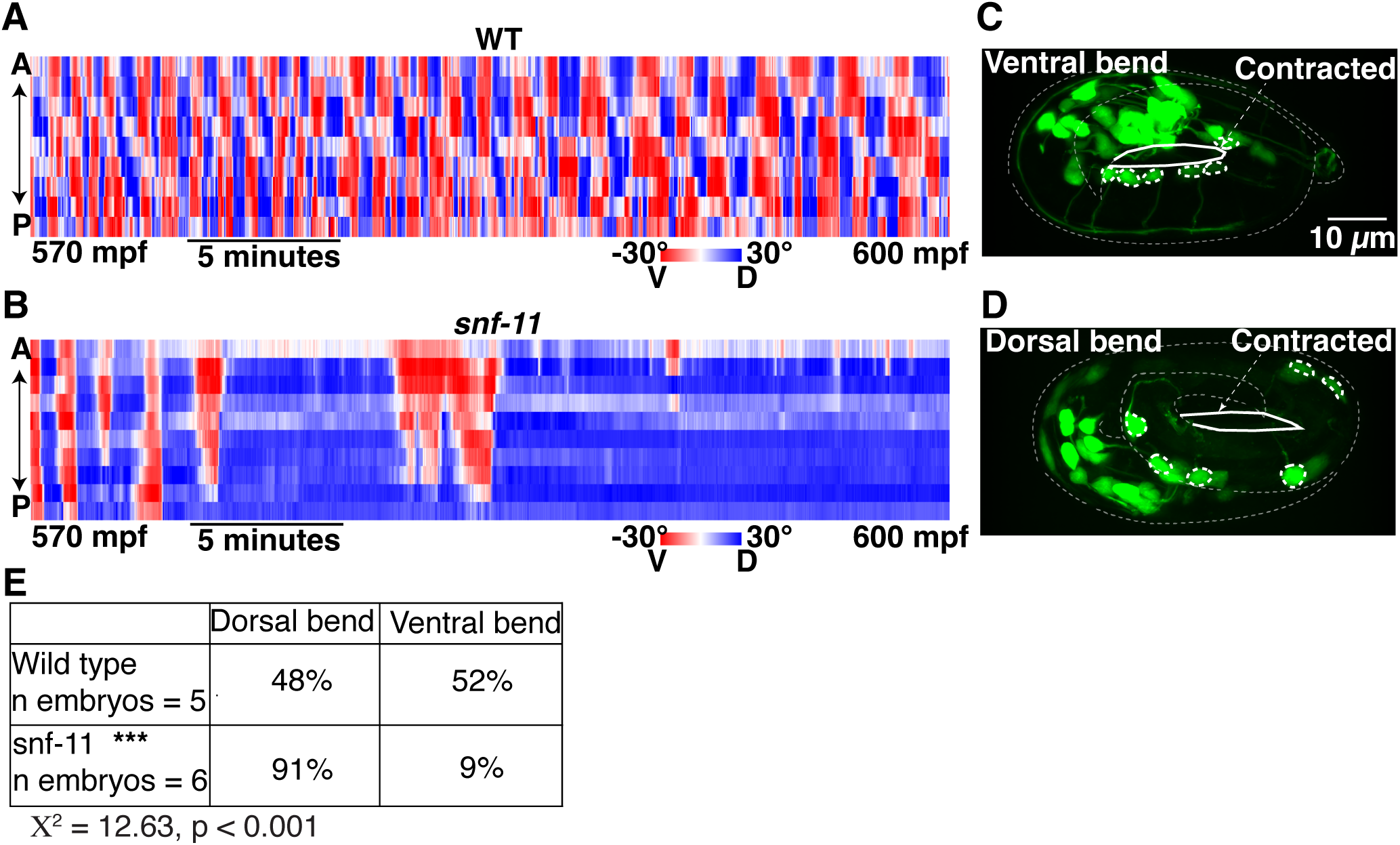
Exaggerated GABA signaling in *snf-11* mutants inhibits flipping behavior. (A-C) Posture analysis is shown for a WT embryo (A) and a *snf-11(nu775)* (B) embryo from 570-600 mpf. Each column is one frame (0.33 seconds) and each row represents dorsoventral bending for each left-right seam cell pair along the anterior-posterior axis. Colors indicate the bend angle (dorsal or ventral) for each seam cell pair. An entirely red column indicates a full body bend in one direction; a full blue column indicates a full body bend in the opposite direction. At this time, WT embryos (A) rapidly alternate between full dorsal and full ventral body coils (i.e. flipping behavior) whereas *snf-11* GAT1 mutants (B) exhibit a prolonged coil in one direction. The WT embryo data (A) was taken from a prior publication (21, 77). (C-E) Comparison of dorsoventral bending in WT and *snf-11* GAT1 mutants. Dorsoventral bending was profiled in embryos expressing soluble GFP in cholinergic neurons (using the *unc-17* promoter). The cholinergic DA/DB motor neuron cell bodies (dashed circles) identify the embryo’s ventral surface. The contracted embryo surface is indicated (solid white outline). Representative images of a dorsally coiled embryo (C), a ventrally coiled embryo (D), and summary data (E) are shown. *snf-11* GAT1 embryos were significantly more likely to exhibit a dorsally coiled posture than WT controls (P<0.001; Χ^2^=12.63).

The embryo posture analysis does not distinguish between dorsal and ventral body bends. Consequently, these data do not indicate if *snf-11* embryos exhibit a dorsal or ventral coiling bias. To address this question, we profiled body bend direction in wild type and *snf-11* embryos at ∼540-600 mpf by confocal microscopy. For this analysis, we used cholinergic (DA/DB) motor neuron cell bodies (expressing GFP) to identify the embryo’s ventral surface. In WT controls, DA/DB cell bodies were equally likely to be on the contracted and relaxed sides of embryo postures (Fig. 3 E-G), consistent with the rapid flipping between dorsal and ventral bends that occurs at this time (21). By contrast, DA/DB cell bodies were significantly more likely to be on the relaxed side of *snf-11* GAT1 mutant embryos (Fig. 3G). We conclude that excess GABA signaling causes embryos to adopt prolonged dorsal body bends. Because *snf-11* mutants exhibited such a strong phenotype, we used them to identify other components required for GABA signaling in early embryos.

### GABA released from DD motor neurons inhibits embryo flipping

We next asked which cells produce the GABA that inhibits embryo motion in *snf-11* GAT1 mutants. *C. elegans* has several classes of GABAergic neurons (41). Different transcription factors specify the differentiation of each class of GABA neurons (42). Embryo motion was restored to wild type levels in *snf-11* double mutants containing an *unc-30* PITX2 mutation, which blocks differentiation of the D-type GABA motor neurons (Fig. 4 A) (43). By contrast, a *lim-6* Lmx1b mutation (which prevents RIS and AVL differentiation) (44) had no effect on the motion of *snf-11* mutants (Fig. 4 A). In addition to D-type GABA neurons, *unc-30* PITX2 is also expressed in DA/DB cholinergic motor neurons in embryos (36). An *unc-3* COE mutation (which blocks DA/DB differentiation) (45) had no effect on the motion of *snf-11* mutant embryos (Fig. 4 B), suggesting that DA/DB defects are unlikely to account for the increased embryo motion seen in *unc-30;snf-11* double mutants. These results suggest that GABA produced by the DD motor neurons (the only D neuron class present in embryos) controls embryo flipping.

**Figure 4.**
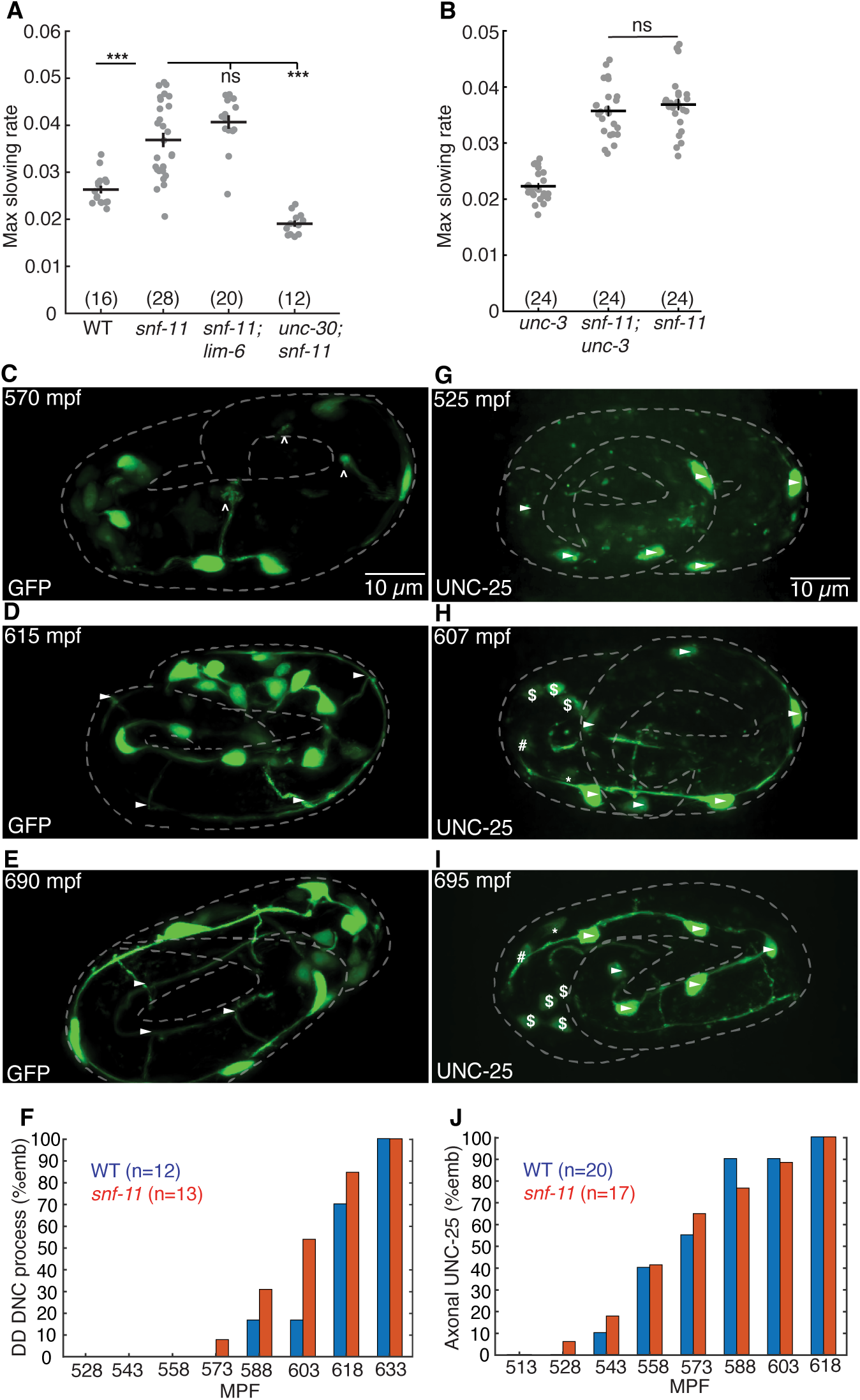
GABA release from immature DD motor neurons inhibits embryo motion. (A-B) Maximal slowing rate of 480-620 mpf embryos are shown for the indicated genotypes.The decreased embryo motion seen in *snf-11(ok156)* GAT1 embryos was unaffected in double mutants lacking *lim-6* Lmx1b (which is required for AVL, RIS, and DVB cell fates) (A), was eliminated in double mutants lacking *unc-30* PITX2 (which is required for the DD cell fate) (A), and was unaffected in double mutants lacking *unc-3* COE (which is required for cholinergic DA/DB motor neuron cell fates) (B). (C-E) Representative images of soluble GFP expressed in DD neurons (using the *unc-47* promoter) is shown at the indicated developmental times. At 570 mpf, DD neurites in the ventral cord are complete but dorsally extending neurites have growth cones and have not yet reached the dorsal cord. At 615 mpf, DD ventral and dorsal neurites, and DD commissures are mostly complete, and little change is observed from 615-690 mpf. Growth cones (carats) and completed commissures (arrowheads) are indicated. (F) Summary data for DD dorsal nerve cord neurite presence over time for wild type (blue, n=12) and *snf-11* (red, n=13) embryos. No significant differences were observed (Chi-squared test; p>0.05). (G-I) Expression of UNC-25(nu808 mNG) is shown at the indicated developmental times. At 525 mpf (G), UNC-25 is observed in six DD cell bodies (arrowheads) but fluorescence is not observed in the DD ventral cord processes. At 607 mpf (H), UNC-25 is observed in DD cell bodies, DD ventral processes, and in some DD dorsal neurites and commissures. At this time, UNC-25 is also seen in 6 head neurons, including RIS (*), AVL (#), and four RME ($) neurons. At 695 mpf (I), UNC-25 is observed in DD cell bodies, DD commissures, and throughout DD ventral and dorsal cord neurites, as well as in the head neurons. (J) Summary data for UNC-25 (nu808 mNG) presence in the ventral process over time for wild type (blue, n=20) and *snf-11* (red, n=17) embryos. No significant differences were observed (Chi-squared test; p>0.05).

Next, we asked if DD neurons are developmentally mature during embryo slowing in *snf-11* GAT1 mutants. DD neurite morphology was examined by expressing GFP in DD neurons (using the *unc-47* VGAT promoter) (Fig. 4 C-F). DD ventral nerve cord processes were complete at 540-570 mpf (Fig. 5 B), while DD commissures and dorsal nerve cord processes were completed after 600 mpf (Fig. 5 D-F). At 570 mpf, DD neurons typically had incomplete commissures with dorsally directed growth cones (Fig. 5 B). We assessed UNC-25 GAD expression using a CRISPR allele that contains mNG inserted into the endogenous *unc-25* locus, just after the ATG codon (Fig. 5 G-J). At 510-540 mpf, UNC-25 (nu808 mNG) appears primarily in the DD motor neurons (Fig. 5 G), whereas expression in other GABA neurons (RIS, AVL, and RME) begins significantly later (after 600 mpf) (Fig. 4 H and I). The onset of UNC-25 (nu808 mNG) expression in DD neurons coincides with the timing of motion inhibition in *snf-11* GAT1 mutants (Fig. 3 A) and the timing of *unc-25* gene transcription in DD neurons (36). Interestingly, UNC-25 (nu808 mNG) was primarily found in DD cell bodies at 510-550 mpf (when motion inhibition occurs in *snf-11* mutants). Very little UNC-25 (nu808 mNG) was detected in DD neuron commissures, ventral cord, and dorsal cord processes until after 560 mpf (Fig. 4 J). Neither the timing of DD neurite outgrowth (Fig. 4 F) nor the appearance of UNC-25(nu808 mNG) in DD neurites (Fig. 4 J) was significantly altered in *snf-11* mutants, indicating that DD maturation was not dramatically altered. These results suggest that immature DD neurons inhibit embryo flipping. These results also suggest that non-synaptic GABA release from DD cell bodies could contribute to flipping inhibition, since UNC-25 GAD protein is restricted to DD cell bodies and DD NMJs are not observed in the ventral cord at this time in development (20).

**Figure 5.**
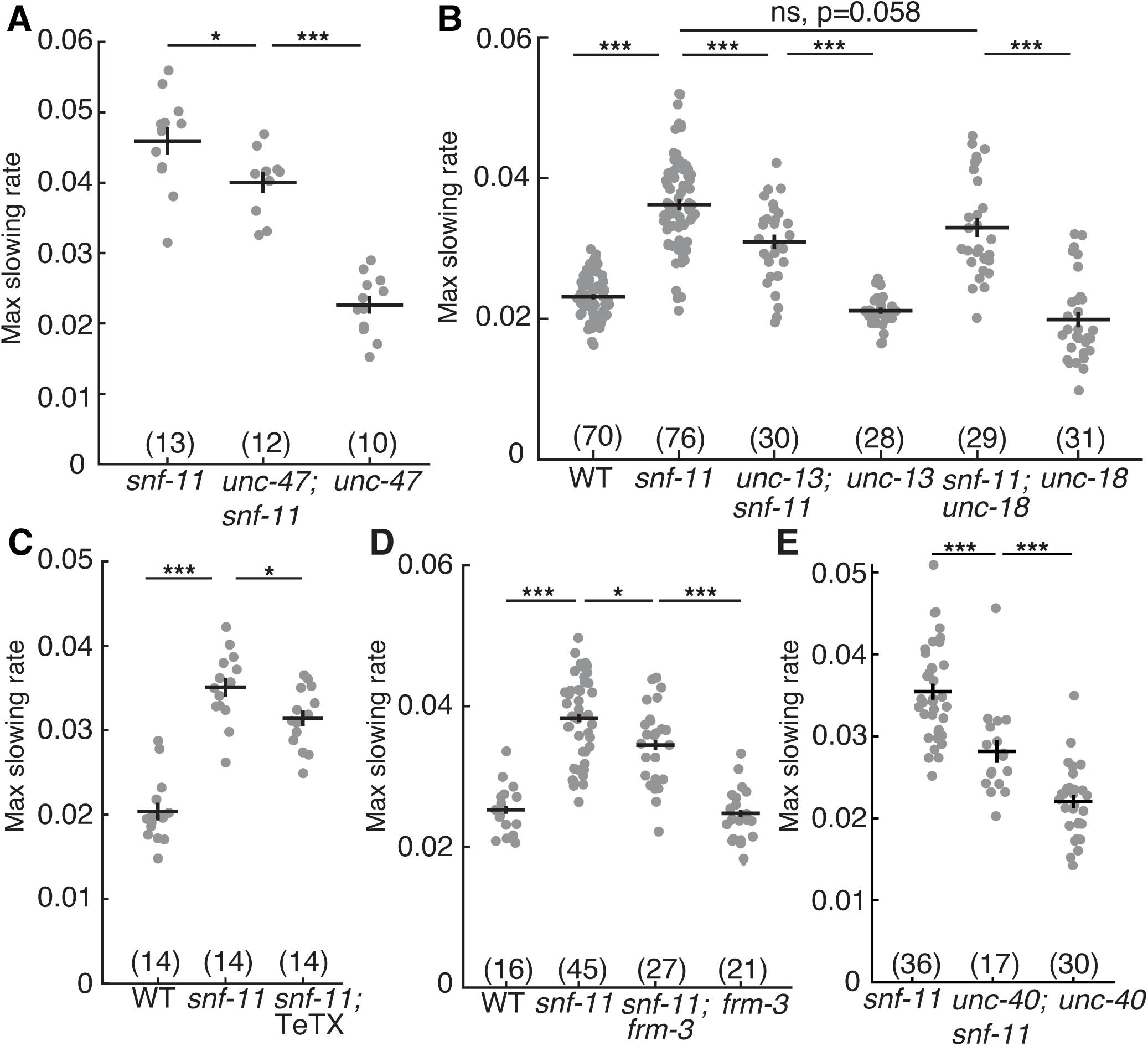
Synaptic and non-synaptic GABA release both contribute to motion inhibition in *snf-11* GAT1 mutants. (A-E) Maximal slowing rates of 480-620 mpf embryos are shown for the indicated genotypes. (A) Inactivating *unc-47* VGAT, which is required to transport GABA into SVs (51), significantly reduces motion inhibition in *snf-11* GAT1 embryos but does not eliminate it. (B) Loss of the SV priming factor *unc-13* MUNC13 partially decreased motion inhibition in *snf-11* GAT1 embryos, while loss of the priming factor *unc-18* MUNC18 did not. (C) GABA neuron expression of tetanus toxin (TeTX), which cleaves the SV SNARE synaptobrevin (53), decreased motion inhibition in *snf-11* GAT1 embryos. (D-E) Two mutations preventing UNC-49 clustering at post-synapses, *frm-3* FARP (D) and *unc-40* DCC (E) decreased motion inhibition in *snf-11(ok156)* (D) and *snf-11(nu775)* (E) mutants. Sample sizes for each genotype are indicated in each figure panel. Values that differ significantly are indicated (ns, not significant; *, *p* <0.05; ***, *p* <0.001). Error bars indicate SEM.

### Muscle UNC-49/GABA_A_ receptors are required for inhibition of flipping

Because DD motor neurons provide synaptic input to body muscles (White 1986), we next asked if muscle GABA receptors control embryo flipping. Consistent with this idea, mutations inactivating UNC-49 GABA_A_ receptors increased embryo motion in *snf-11* double mutants and this effect was reversed by a transgene that restores UNC-49 expression in body muscles (*SI Appendix,* Fig. S3A). By contrast, inactivating subunits that form the GABA activated G protein coupled receptor (GBB-1 and GBB-2) had no effect on *snf-11* embryo motion (*SI Appendix,* Fig. S3B). Collectively, these results suggest that inhibition of embryo flipping in *snf-11* GAT1 mutants is mediated by GABA released by the DD motor neurons, which activates UNC-49 GABA_A_ receptors in body muscles.

### GABA inhibits embryo flipping by hyperpolarizing body muscles

In mammals, GABA often has excitatory effects on immature neurons and only becomes inhibitory as neurons mature (46, 47). This pattern of early excitatory GABA effects is also thought to occur in *C. elegans* (48). Prompted by these results, we next asked if GABA’s effects on embryo motion are mediated by depolarizing or hyperpolarizing body muscles. Because UNC-49 GABA_A_ subunits form GABA-activated chloride channels, GABA’s impact on muscle activity depends on the transmembrane chloride gradient. UNC-49 activation hyperpolarizes muscles when extracellular chloride exceeds cytoplasmic levels, while UNC-49 depolarizes muscles in the converse conditions. The transmembrane chloride gradient is controlled by the activity of two chloride ion extruders (ABTS-1 and KCC-2) and one chloride ion importer (NKCC-1) (48, 49). Loss of the chloride importer NKCC-1 further inhibits motion in *snf-11* double mutants, loss of the chloride extruder KCC-2 increased motion in *snf-11* double mutants, while loss of the ABTS-1 chloride extruder had no effect (*SI Appendix,* Fig. S3 C-E). Taken together, these results suggest that GABA prolongs dorsally coiled postures by hyperpolarizing ventral body muscles. GABA’s preferential effect on ventral body muscles most likely results from the fact that UNC-49 receptors are found in ventral but not dorsal body muscles in both embryos (*SI Appendix,* Fig. S3 F) and newly hatched first stage larvae (50).

### SV exocytosis modestly contributes to GABA mediated inhibition of embryo motion

We next asked if SNF-11’s impact on embryo motion is mediated by synaptic GABA release. Synaptic GABA release is mediated by exocytosis of SVs at pre-synaptic active zones. To test its role in controlling early embryo motion, we constructed *snf-11* GAT1 double mutants containing mutations that prevent synaptic GABA release. Embryo motion in *snf-11* GAT1 mutants was modestly increased by: mutations that prevent SV loading with GABA (*unc-47* VGAT) (51) (Fig. 5 A) or SV docking (*unc-13* MUNC13) (52) (Fig. 5 B); and by expression of tetanus toxin in DD neurons, which cleaves the vesicle SNARE protein synaptobrevin (53) (Fig. 5 C). In all cases, *snf-11* GAT1 double mutants lacking synaptic GABA release had significantly less motion than that observed in *unc-25* GAD; *snf-11* GAT1 double mutants (Fig. 2 F), suggesting that DD neurons exhibit significant residual GABA release even when synaptic release was blocked. To further address the role of vesicular GABA release, we also analyzed *unc-18* mutants (which reduce SV exocytosis) (54) and *cat-1* VMAT mutants, because VMAT2 promotes GABA transport into SVs in mouse dopaminergic neurons (55). Embryo motion in *snf-11* GAT1 mutants was not affected by *cat-1* VMAT mutations (*SI Appendix,* Fig. S4A). There was a trend for increased motion in *snf-11; unc-18* double mutants, which was not statistically significant (*p*= 0.058, Fig. 5 B). Finally, it is possible that synaptic GABA release inhibits embryo motion only in *snf-11* GAT1 mutants, where GABA signaling is exaggerated. Contrary to this idea, several mutations that impair synaptic GABA release significantly increased embryo motion in single mutants (*SI Appendix,* Fig. S4B). Taken together, these results suggest that synaptic release contributes to GABA mediated inhibition of embryo flipping.

### Synaptic UNC-49/GABA_A_ receptors contribute to GABA mediated inhibition of embryo flipping

Thus far, our results suggest that SV exocytosis contributes to GABA release from DD neurons in early embryos (510-570 mpf). In mature neurons, SV exocytosis is restricted to pre-synaptic active zones; however, SV exocytosis could occur at non-synaptic sites in immature neurons. To further address whether GABA is released at synapses, we asked if clustered post-synaptic GABA receptors are required for inhibiting embryo motion. To address this possibility, we analyzed mutants lacking two scaffolding proteins (FRM-3/FARP and UNC-40/DCC) that immobilize UNC-49 GABA_A_ receptors at post-synaptic elements (56, 57). We found that embryo motion was modestly increased in both *frm-3* FARP; *snf-11* GAT1 and *unc-40* DCC; *snf-11* GAT1 double mutants compared to *snf-11* single mutant controls (Fig. 5 D and E). The magnitude of these increases in embryo motion are similar to those in *snf-11* GAT1 double mutants lacking UNC-47 VGAT or UNC-13. These results suggest that synaptic GABA release and activation of clustered synaptic UNC-49 GABA_A_ receptors both contribute to the inhibited embryo motion seen in *snf-11* GAT1 mutants.

### Bestrophin channels promote GABA mediated inhibition of embryo motion

Because mutations that eliminate synaptic GABA transmission only partially block inhibition of embryo flipping in *snf-11* GAT1 mutants, we next asked if immature DD neurons also release GABA by a non-synaptic mechanism. In tonic transmission, GABA is released by plasma membrane channels rather than by SV exocytosis (58). In mammals, two classes of channels are implicated in tonic GABA release: bestrophin channels and SWELL1 (LRC88A) osmolyte channels (58). The *C. elegans* genome encodes 26 bestrophin paralogs but lacks an obvious SWELL1 equivalent. Two bestrophin genes (*best-18* and *best-19*) are weakly expressed in embryonic DD motor neurons (36). We therefore asked if *best-18* or *best-19* are required for GABA to inhibit motion in *snf-11* GAT1 mutant embryos. Embryo motion was significantly increased in *best-18; best-19; snf-11* triple mutants as well as in *best-18; snf-11* double mutants (compared to *snf-11* single mutants) (Fig. 6A). The motion of *best-18; best-19; snf-11* triple mutant embryos was significantly decreased by transgenes expressing *best-19* in DD neurons using either of two promoters (*unc-30* or *cpg-8*) (*SI Appendix,* Fig. S5A), suggesting that BEST-19 functions in DD neurons to promote GABA signaling. The *unc-25* GAD mutant had significantly more embryo motion than in *best-18; best-19; snf-11* triple mutants (Fig. 6A), suggesting that DD neurons had significant residual GABA release in the triple mutants.

**Figure 6.**
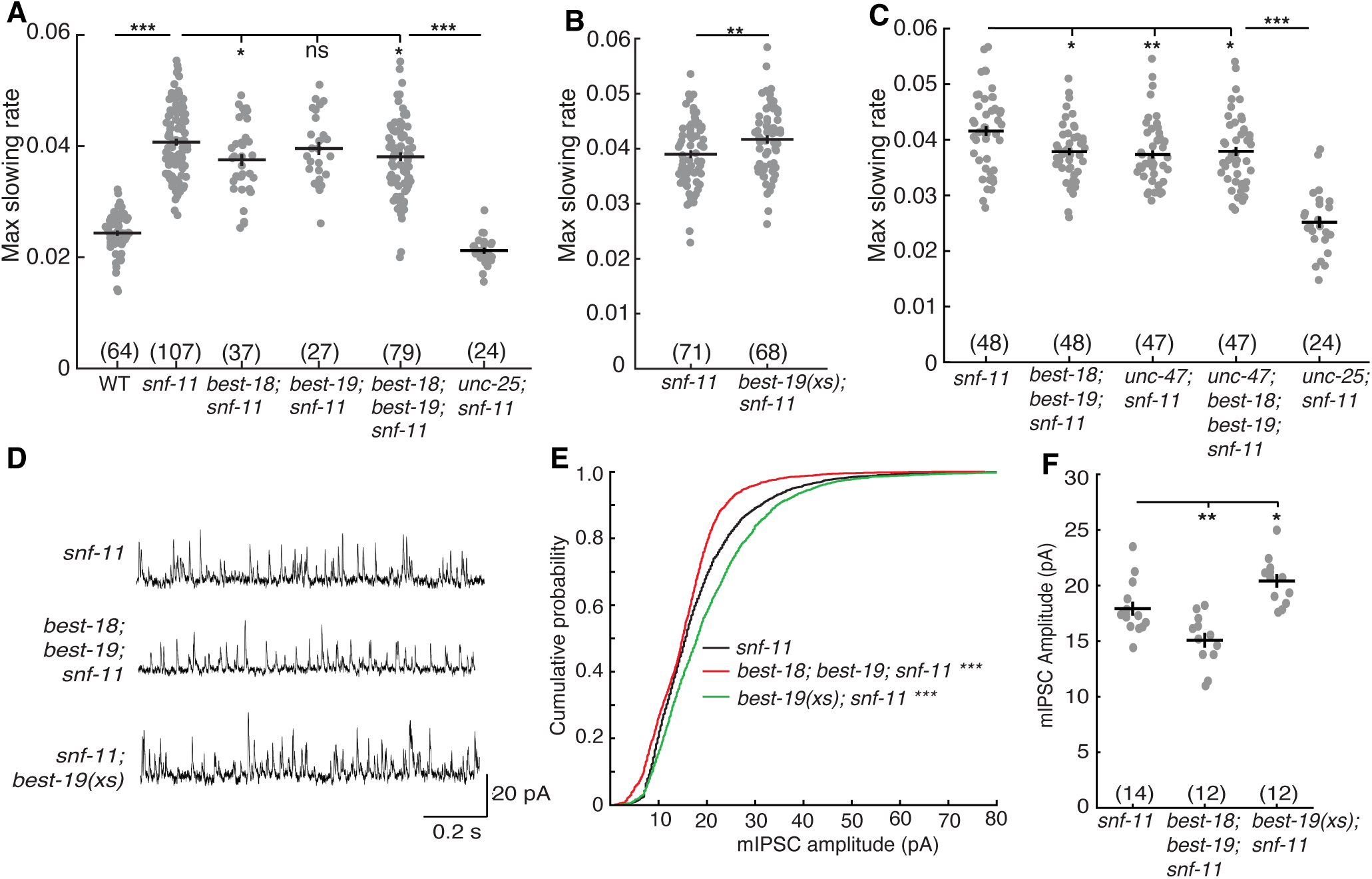
Bestrophins promote synaptic GABA transmission. (A-C) Maximal slowing rate of 480-620 mpf embryos is plotted for the indicated genotypes. (A) Inhibition of embryo motion in *snf-11* GAT1 mutants is modestly decreased in mutants lacking BEST-18, and in double mutants lacking both BEST-18 and BEST-19 but is not significantly altered in mutants lacking just BEST-19. (B) BEST-19 over-expression in GABA neurons further inhibits embryo motion in *snf-11* mutants. (C) Inactivating *unc-47* VGAT and bestrophins do not have additive effects on embryo motion in *snf-11* mutants, implying that UNC-47 and Bestrophins act together to inhibit embryo motion.(D-F) Bestrophin inactivation and over expression alter synaptic transmission at adult DD NMJs. Representative mIPSC traces from adult muscles (D), cumulative probability distrbutions for mIPSC amplitude (E), and mean mIPSC amplitudes (F) are shown for the indicated genotypes. mIPSC amplitudes are significantly increased in *snf-11; best-19(xs)* and decreased in *best-18; best-19; snf-11* triple mutants. Sample sizes for each genotype are indicated in each figure panel. Values that differ significantly are indicated (ns, not significant; *, *p* <0.05; **, *p* <0.01; ***, *p* <0.001). Error bars indicate SEM.

Bestrophin mediated GABA release has been documented in glia in both *C. elegans* and mammals (59, 60) but has not been described in neurons. To further test their role in GABA release from neurons, we asked if increased Bestrophin gene expression could further reduce flipping in *snf-11* embryos. Consistent with this idea, overexpressing BEST-19 in GABA neurons (using the *unc-25* GAD promoter) inhibits motion in *snf-11* GAT1 mutant embryos (Fig. 6B). Thus, inactivating BEST-18 and BEST-19 increased embryo motion while over-expressing BEST-19 in GABA neurons decreased motion. By contrast, loss and gain of bestrophin function had no effect in *unc-25; snf-11* embryos, indicating bestrophins likely act via GABA signaling (*SI Appendix,* Fig. S5B).

### Bestrophin channels promote synaptic GABA transmission

If bestrophin channels mediate non-synaptic GABA release, bestrophin mutations and mutations inactivating synaptic GABA release should have additive effects on embryo behavior. However, we found no significant differences in motion inhibition between *unc-47; snf-11* double mutants (which lack GABA SV loading), *best-18; best-19; snf-11* triple mutants, and *best-18; unc-47; best-19; snf-11* quadruple mutants (Fig. 6C). We further find that bestrophins and *unc-13* (which lacks SV exocytosis) also do not have additive effects on embryo behavior; in fact, *unc-13; best-18; best-19; snf-11* embryos have significantly more motion inhibition than *unc-13; snf-11* embryos (*SI Appendix,* Fig. S5C). By contrast, embryo motion of *unc-13; snf-11* double mutants and *unc-13; unc-47; snf-11* triple mutants were not significantly different (*SI Appendix,* Fig. S5D), consistent with UNC-13 and UNC-47 acting together to promote synaptic GABA release. Lack of additivity with *unc-47* VGAT and *unc-13* mutations suggests that BEST-18 and BEST-19 promote synaptic GABA release.

One possible explanation for these results is that bestrophins are required for UNC-47/VGAT mediated GABA transport into synaptic vesicles. To test this idea, we recorded miniature inhibitory post-synaptic currents (mIPSCs), which corresponds to the synaptic current evoked by fusion of a single SV. Thus, mIPSC amplitude scales with the amount of GABA packaged into SVs. In adult body muscles, mIPSC amplitudes were significantly decreased in double mutants lacking BEST-18 and BEST-19 and were significantly increased in transgenic animals overexpressing BEST-19 in GABA neurons (Fig. 6D-F). These results provide further support for the idea that bestrophins promote synaptic GABA signaling.

How do bestrophins alter mIPSC amplitude? A chloride channel (CLC-3) promotes acidification of SVs, thereby promoting VGAT mediated GABA transport into SVs (61). To determine if bestrophins (which are also permeable to chloride ions) (62) promote SV acidification, we analyzed synaptopHluorin (SpH) fluorescence at DD synapses (*SI Appendix,* Fig. S6A-C). SpH contains a pH-sensitive GFP variant fused to the lumenal domain of SNB-1/synaptobrevin (63, 64). We expressed SpH carrying an N-terminal scarlet tag in GABA neurons (using the *unc-25* promoter) (*SI Appendix,* Fig. S6A and B). SpH was restricted to the intracellular SV pool by analyzing Scarlet-SpH fluorescence in *unc-13* mutants, which lack a surface SpH pool (65). To assess SV acidification, we measured the Green/Red fluorescence ratio (G/R) produced by Scarlet-SpH. Scarlet-SpH G/R ratios were not significantly altered in *best-18; best-19* double mutants nor in transgenic animals over-expressing BEST-19 in GABA neurons (*SI Appendix,* Fig. S6C), suggesting that the lumenal pH of SVs had not been significantly altered.

An alternative explanation for changes in mIPSC amplitude is that bestrophins alter UNC-49/GABA_A_ abundance at post-synaptic elements. Contrary to this idea, endogenous UNC-49 levels at adult GABAergic neuromuscular junctions were not detectably altered in *best-18; best-19; snf-11* triple mutants nor in adults over-expressing BEST-19 in GABA neurons but were significantly reduced in *frm-3* FARP mutants (*SI Appendix,* Fig. S6 D and E). These results suggest that changes in mIPSC amplitudes are unlikely to result from a change in synaptic UNC-49 levels. Taken together, these results suggest that bestrophin channels promote synaptic GABA release from DD motor neurons, most likely by enhancing GABA loading into SVs and not by acting as plasma membrane GABA channels, as in glia.

## Discussion

Our results lead to five principal conclusions. First, most nerve ring synapses form after 600 mpf, which coincides with the onset of UNC-13 dependent locomotion behavior. Second, embryo flipping behavior is inhibited by GABA release from immature DD motor neurons. Third, GABA inhibits embryo flipping by hyperpolarizing ventral body muscles. Fourth, DD neurons inhibit embryo flipping via both synaptic and non-synaptic (i.e. tonic) GABA release. Fifth, bestrophins function in DD neurons to promote synaptic GABA transmission. Below we discuss the significance of these findings.

### Most nerve ring synapses form after 600 mpf

When do synapses first form and exert control over embryo behavior? Prior studies gave conflicting answers to this question. Several active zone proteins (including SYD-2/liprin-a, UNC-10/RIM, ELKS-1, CLA-1/Clarinet) were reported to be present in nerve ring axons at ∼ 420 mpf (19). Small NMJs were seen in electron micrographs of the dorsal nerve cord of 550 mpf embryos, whereas ventral NMJs were not seen at this time (20). Mutations inactivating UNC-13 strongly disrupt sinusoidal locomotion and rhythmic quiescent bouts at 650 mpf, but did not alter motion prior to 600 mpf (21). Thus, analysis of synapse structure (using active zone proteins or electron micrographs) and function (using behavior) provide different estimates for when synapses form during embryogenesis. To reconcile the differences between these prior studies, we analyzed localization of 3 synaptic ion channels (UNC-2/CaV2, UNC-49/GABA_A_, and ACR-16/CHRNA7) and found that all 3 appear in the nerve ring after 600 mpf, which was ∼1-3 hours after the onset of their mRNA expression. These channels are essential for synaptic function; consequently, their nerve ring localization provides a good estimate for when these synapses begin to function. It is noteworthy that the nerve ring localization of these ion channels coincides with the timing of UNC-13 dependent behaviors (21). These results suggest that most nerve ring synapses form after 600 mpf and that earlier arriving active zone proteins represent intermediates in synapse assembly. These results further suggest that non-synaptic signaling mechanisms are likely to be involved in behaviors occurring before 600 mpf. These results do not exclude the possibility that these ion channels are present at low levels at some nerve ring synapses prior to 600 mpf, since low levels of mNG fluorescence may not be detected. Finally, these results highlight the importance of combining structural and functional studies to analyze synapse development.

### GABA release from immature DD neurons inhibits embryo flipping

Using embryo motion to assess nervous system function, we identify an early role for GABA release inhibiting alternation of full body bends (flipping). Flipping is strongly inhibited in embryos with exaggerated GABA signaling (i.e. in *snf-11* GAT1 mutants), and inhibition of flipping is eliminated by mutations inactivating the GABA biosynthetic enzyme UNC-25 GAD, UNC-49 GABA_A_ receptor expression in body muscles, and UNC-30 PITX2 (which is required for DD neuron differentiation). Flipping inhibition appears at approximately 510-570 mpf, before DD neuron commissures and dorsal cord neurites are completed, before UNC-25 GAD is observed in DD ventral cord processes, and before DD NMJs are observed in the ventral cord (20). Thus, immature DD neurons inhibit embryo flipping and this represents a very early neuronally controlled behavior, prior to formation of most nerve ring synapses.

### Components of the SV release machinery contribute to but are not essential for GABA release in early embryos

Synaptic neurotransmitter release is crucial for adult *C.* elegans movement and viability (66, 67). In late embryos (650 mpf), a strong *unc-13* loss of function mutation disrupts sinusoidal motion and rhythmic behavioral quiescence (21). In early embryos, we find that loss of GABA synthesis eliminates the inhibition of embryo motion in *snf-11* GAT1 mutants. However, mutations preventing GABA loading into SVs (*unc-47* VGAT), those preventing SV fusion (*unc-13* and tetanus toxin expression), and those preventing post-synaptic clustering of UNC-49 GABA_A_ (*frm-3* FARP and *unc-40* DCC) only modestly (but significantly) increased embryo motion in *snf-11* GAT1 mutants. These results suggest that synaptic GABA transmission commences by 510-570 mpf, before DD neurite outgrowth is completed, and before DD neuron NMJs are observed in the ventral cord (20). These results also suggest that non-synaptic GABA release from DD neurons plays a significant role in inhibition of embryo flipping behavior. Further experiments will be required to determine the mechanism mediating non-synaptic GABA release.

### Bestrophin channels promote synaptic GABA release

Bestrophin channels (plasma membrane channels that flux chloride, GABA, and glutamate) mediate non-synaptic tonic GABA release from glia (68). Surprisingly, we find that bestrophin channels contribute to synaptic GABA release from DD motor neurons. Several results support this conclusion. First, BEST-19 over-expression in GABA neurons significantly enhanced embryo slowing in *snf-11* mutants. Second, expression of BEST-19 in DD neurons rescued the embryo slowing defect in *best-18; best-19;snf-11* triple mutants. Third, *best-18* and *best-19* mutations and mutations that prevent synaptic GABA release (*unc-47* VGAT and *unc-13* MUNC13) did not have additive effects on GABA mediated inhibition of embryo motion, suggesting that bestrophins and UNC-47 function together to promote synaptic GABA transmission. Fourth, decreased and increased bestrophin function in GABA neurons had reciprocal effects on adult mIPSC amplitudes, indicating that bestrophins promote synaptic GABA transmission. Taken together, these results strongly support the idea that BEST-18 and BEST-19 act in DD neurons to promote synaptic GABA signaling.

How do bestrophins promote synaptic GABA transmission? We can imagine three potential mechanisms. First, if BEST-18 and BEST-19 bind GAD (as previously reported for mammalian BEST1) (69), they could function as scaffold proteins that recruit UNC-25/GAD to pre-synaptic elements, thereby promoting GABA loading into SVs. Second, BEST-18 and BEST-19 channels (which like all known bestrophins are likely permeable to chloride ions) (62) could promote SV acidification, as previously shown for mammalian CLC-3 chloride channels (61). Contrary, to this hypothesis, bestrophin mutations and over-expression did not alter SV acidification. Third, BEST-18 and BEST-19 could increase UNC-49 levels at DD NMJs; however, bestrophin mutations and over-expression did not alter post-synaptic UNC-49 levels in the dorsal nerve cord. Our results do not exclude any of these hypotheses because we could have missed small changes in SV pH or in synaptic UNC-49 levels. Thus, further experiments will be required to determine how BEST-18 and BEST-19 promote synaptic GABA transmission. Because BEST1 is expressed in mouse cortical neurons (70), bestrophins could also promote synaptic GABA transmission in vertebrates.

Loss of bestrophins caused a minor reduction in mIPSC amplitude, while loss of *unc-47*/VGAT nearly abolishes mIPSCs (71). Yet, loss of bestrophins and loss of *unc-47*/VGAT had similar effects on behavior in *snf-11* mutant embryos. It is possible that bestrophins have a more significant impact on GABA vesicle loading in early embryos, when GAD expression has only just begun, GAD is primarily retained in DD cell bodies, and DD synapse formation is incomplete.

### ASD risk genes function in GABA signaling pathways active in early embryonic development, a critical period for ASD

Multiple genes involved in GABA transmission are linked to neurodevelopmental disorders such as ASD (27–29). Understanding the functions of ASD risk genes and the pathways in which they function during embryonic development is of particular importance. Here we show that multiple genes associated with ASD, including *snf-11* GAT1 and *unc-49* GABA_A_R, function in early embryo GABA signaling to regulate behavior. In *C elegans*, UNC-49 serves as the GABA_A_ receptor at both synaptic and extra-synaptic sites in body muscles. By contrast, vertebrate synaptic and extra-synaptic GABA_A_ receptors have distinct subunits, which are encoded by different genes (72). Additionally, our results raise the possibility that early effects of GABA on nervous system development and ASD could be mediated by non-synaptic modes of transmission. Further research on the early functions of *snf-11* GAT1 and GABA could provide insight into how GABA pathway mutations confer risk for ASD and other neuropsychiatric disorders. Similarly, it is possible that non-synaptic signaling by other neurotransmitters (i.e. in addition to GABA) also plays an important role in shaping early brain activity.

## Materials and Methods

Detailed information on strains, reagents, experimental protocols, and data analysis can be found in the Extended Methods (Supporting Information).

### Strains and Reagents

Animals were maintained on nematode growth medium (NGM) seeded with *Escherichia coli* (OP50) at room temperature as described in Brenner (1974) (73).

### Brightfield embryo motion assay

Assays were performed as described in Ardiel et al. (2022) (21). Briefly, embryos were dissected from gravid day 1 adults and arrayed on poly-L-lysine (0.1 mg/mL) in an M9 filled glass bottom dish (MatTek Corp., P35G-1.5–20 C). Embryos were then imaged at 1 Hz on an inverted microscope (Zeiss, Axiovert 100) at room temperature.

To quantify embryo motion, pixel intensities were collected from a box within each embryo. Frame-to-frame pixel intensity changes larger than a 100 AU threshold were counted. Embryonic age was determined by measuring time following twitch onset, which was defined as 430 mpf. Minutes post fertilization values for developmental landmarks (such as hatch time) exhibit some day-to-day variation, most likely due to subtle differences in ambient temperature.

### Inhibition of embryo motion was quantified as follows

1. For each embryo, 2,500 second (41.67 minute) twitch profiles were extracted with a sliding window (200 s step size) from 60 minutes post twitch until 150 minutes post twitch. Missing datapoints were replaced with a reversed duplication of the immediately preceding interval of the same length.
2. Using MATLAB’s polyfit function, the regressed linear slope was calculated for each 2,500 s twitch profile.
3. For each embryo, the timing and magnitude of the most negative slope (pixels changing over threshold/s) was recorded.
4. One-way ANOVA (implemented in Matlab) was used to statistically test whether strains differed in minimum slope. A post-hoc Tukey-Kramer test was applied to correct for multiple comparisons.

### diSpim imaging and posture analysis

diSpim imaging was performed as described in Ardiel et al. (2022) (21) from ∼90 to 180 minutes after twitch onset, as estimated based on observation of embryos under a dissecting microscope. Embryos were dissected as described above. On a diSPIM (74), a pair of perpendicular water-dipping, long-working distance objectives (40 x, 0.8 NA) were used for brightfield and fluorescence imaging. Acquisition was controlled using Micro-Manager’s diSPIM plugin (https://micro-manager.org/). Imaging was done in iSPIM mode, using a single view. Volumes were acquired at 3 Hz. Detailed protocols for diSPIM embryo imaging can be found in Duncan et al. (2019) (75).

*Snf-11* postures were built using *wIs51[SCM*p::GFP + *unc-119*(+)], which expresses GFP in hypodermal seam cell nuclei. Seam cell fluorescence produced by the *wIs51* transgene was too dim to accurately track postures before ∼570 mpf. Images were segmented as described in detail in Ardiel et al., 2022 (21). Seam cells were manually positioned along the body axis by observation of several frames to produce a seed volume. From there, seam cell tracking was performed using a Global Nearest Neighbor (GNN) algorithm as implemented in Matlab. Cell identifications were manually corrected when tracking errors became apparent. Dorsoventral bends were computed from the midpoint along the spline connecting neighboring seam cells and the corresponding midpoint on the opposite side of the body. See Ardiel et al., 2022 (21) for details.

### Fluorescence imaging of embryos

Embryos were dissected from gravid day 1 adults as described under “Brightfield Motion Assay” and synchronized at the 2-cell stage. Embryos were then moved to an inverted confocal microscope (Nikon, Eclipse Ti2) and repeatedly imaged using a 60X/1.49NA objective using the resonant scanner. All images were acquired at room temperature. Image denoising was performed using the Denoise.AI algorithm (Nikon). All subsequent image analyses were done using FIJI.

### Nerve ring synapse imaging

Synaptic ion channel and adhesion molecule abundance in the nerve ring was assessed by quantifying integrated nerve ring fluorescence as follows: images were thresholded at 40 AU, the nerve ring region of interest was identified by manual inspection, and the total fluorescence within the thresholded volume was measured using the FIJI 3D Objects Counter function.

### Dorsoventral bend bias assay

Dorsal or ventral bend was manually assessed based on the vsIs48 [*Punc-17:GFP*] marker, which labels cholinergic neurons in the head and along the ventral cord. Differences between strains were assessed using the Chi-square test.

### Dorsoventral UNC-49 expression assay

Expression of UNC-49(nu829 mNG) in ventral and dorsal muscles of the embryo was manually assessed using the otIs374[*Punc-47:mChopti*] to label GABA neuron cell bodies on ventral side of the embryo.

### DD neurite morphology assay

DD neurite outgrowth was manually assessed based on the oxIs12[*Punc-47:GFP]* marker, which labels GABA neurons.

### DD UNC-25 localization assay

Cell identification and subcellular localization were manually assessed in confocal images based on UNC-25(nu808 mNG) fluorescence.

### Muscle mIPSC recordings

Whole-cell patch-clamp measurements were performed using an Axopatch 200B amplifier with pClamp 10 software (Molecular Devices). The data were sampled at 10 kHz and filtered at 5 kHz. Body muscle IPSCs were recorded as previously described (56). Dissected adults were superfused in an extracellular solution containing 127 mM NaCl, 5 mM KCl, 26 mM NaHCO_3_, 1.25 mM NaH_2_PO_4_, 10 mM glucose, 5mM sucrose, 1 mM CaCl_2_, and 4 mM MgCl_2_, bubbled with 5% CO_2_, 95% O_2_ at 22°C. The pipette solution contained 105 mM CH_3_O_3_SCs, 10 mM CsCl, 15 mM CsF, 4mM MgCl_2_, 5mM EGTA, 0.25mM CaCl_2_, 10mM HEPES, and 5mM Na_2_ATP, 1mM Na_2_GTP adjusted to pH 7.2 using CsOH. Whole-cell recordings were carried out at 0mV to record mIPSCs.

### Fluorescence imaging of adults

Day 1 adult worms were immobilized and the dorsal nerve cord just anterior to the vulva was imaged. Maximum intensity projections for each volume were auto-thresholded and puncta were identified as round fluorescent objects (area >0.1 μm^2^) using analysis of particles. Mean fluorescent intensity in each punctum was analyzed in the raw images.

### UNC-49 puncta analysis

Pre-synaptic regions of interest (ROIs) were identified by localization of an mCherry-tagged synaptic vesicle marker (UNC-57 Endophilin) expressed in the GABA neurons. The intensity of UNC-49(nu829 mNG) in the UNC-57 ROIs was quantified. All image analysis was done using FIJI.

### Synaptic vesicle acidification assay

Pre-synaptic regions of interest (ROIs) in adult dorsal cords were identified by localization of a BFP-tagged active zone marker (ELKS-1 ERC) expressed in the GABA neurons. SV acidification was assessed using a transgene containing SNB-1 Synaptobrevin tagged with Scarleti3 at the N-terminus and super ecliptic pHluorin at the C-terminus (Scarlet-SpH), expressed in GABA neurons. Scarlet-SpH was analyzed in *unc-13* mutants to restrict SpH to the intracellular SV pool (65). SV acidification was assessed by measuring the Scarlet-SpH Green/Red fluorescence ratio at dorsal cord DD synapses in adults. All image analysis was done using FIJI.

## Acknowledgments

We thank the following for strains, advice, reagents, and comments on the manuscript: *C. elegans* genetics stock center (CGC), Shohei Mitani, and members of the Kaplan lab. We thank Abhishek Kumar and the Woods Hole Marine Biological Laboratory imaging center for assistance with diSPIM imaging. This work was supported by an NIH research grant to J.K. (NS32196). The CGC is funded by the National Institutes of Health Office of Research Infrastructure Programs (P40 OD010440).

## Data availability

All Data reported in this paper are included in the Supplemental Appendix.

## References

1. T. K. Hensch, Critical period regulation. Annu Rev Neurosci 27, 549–579 (2004).

2. K. Krishnan et al., MeCP2 regulates the timing of critical period plasticity that shapes functional connectivity in primary visual cortex. Proc Natl Acad Sci U S A 112, E4782–4791 (2015).

3. J. Bock, K. Braun, Blockade of N-methyl-D-aspartate receptor activation suppresses learning-induced synaptic elimination. Proc Natl Acad Sci U S A 96, 2485–2490 (1999).

4. M. K. Mwaniki, M. Atieno, J. E. Lawn, C. R. Newton, Long-term neurodevelopmental outcomes after intrauterine and neonatal insults: a systematic review. Lancet 379, 445–452 (2012).

5. S. Saigal, L. W. Doyle, An overview of mortality and sequelae of preterm birth from infancy to adulthood. Lancet 371, 261–269 (2008).

6. S. N. Mattson et al., Further development of a neurobehavioral profile of fetal alcohol spectrum disorders. Alcohol Clin Exp Res 37, 517–528 (2013).

7. J. E. Sulston, E. Schierenberg, J. G. White, J. N. Thomson, The embryonic cell lineage of the nematode *Caenorhabditis elegans*. Dev. Biol. 100, 64–119 (1983).

8. A. Carreira-Rosario, R. A. York, M. Choi, C. Q. Doe, T. R. Clandinin, Mechanosensory input during circuit formation shapes Drosophila motor behavior through patterned spontaneous network activity. Curr Biol 31, 5341–5349 e5344 (2021).

9. V. Hamburger, Some Aspects of the Embryology of Behavior. Q Rev Biol 38, 342–365 (1963).

10. L. A. Kirkby, G. S. Sack, A. Firl, M. B. Feller, A role for correlated spontaneous activity in the assembly of neural circuits. Neuron 80, 1129–1144 (2013).

11. M. Fagiolini, T. K. Hensch, Inhibitory threshold for critical-period activation in primary visual cortex. Nature 404, 183–186 (2000).

12. B. Williams, R. Waterston, Genes critical for muscle development and function in Caenorhabditis elegans identified through lethal mutations. J. of Cell Biol. 124, 475–490 (1994).

13. S. H. Young, M. M. Poo, Spontaneous release of transmitter from growth cones of embryonic neurones. Nature 305, 634–637 (1983).

14. R. I. Hume, L. W. Role, G. D. Fischbach, Acetylcholine release from growth cones detected with patches of acetylcholine receptor-rich membranes. Nature 305, 632–634 (1983).

15. W. Lin et al., Aberrant development of motor axons and neuromuscular synapses in erbB2-deficient mice. Proc Natl Acad Sci U S A 97, 1299–1304 (2000).

16. M. Munz et al., Pyramidal neurons form active, transient, multilayered circuits perturbed by autism-associated mutations at the inception of neocortex. Cell 186, 1930–1949 e1931 (2023).

17. M. Demarque et al., Paracrine intercellular communication by a Ca2+- and SNARE-independent release of GABA and glutamate prior to synapse formation. Neuron 36, 1051–1061 (2002).

18. M. W. Moyle et al., Structural and developmental principles of neuropil assembly in C. elegans. bioRxiv 10.1101/2020.03.15.992222, 2020.2003.2015.992222 (2020).

19. N. A. McDonald, R. D. Fetter, K. Shen, Assembly of synaptic active zones requires phase separation of scaffold molecules. Nature 588, 454–458 (2020).

20. R. Durbin (1987) Studies on the development and organisation of the nervous system of C. elegans. (University of Cambridge).

21. E. L. Ardiel et al., Stereotyped behavioral maturation and rhythmic quiescence in C. elegans embryos. Elife 11 (2022).

22. J. E. Richmond, W. S. Davis, E. M. Jorgensen, UNC-13 is required for synaptic vesicle fusion in C. elegans. Nat Neurosci 2, 959–964 (1999).

23. C. J. Akerman, H. T. Cline, Refining the roles of GABAergic signaling during neural circuit formation. Trends Neurosci 30, 382–389 (2007).

24. Z. J. Huang, Activity-dependent development of inhibitory synapses and innervation pattern: role of GABA signalling and beyond. J Physiol 587, 1881–1888 (2009).

25. Z. J. Huang, P. Scheiffele, GABA and neuroligin signaling: linking synaptic activity and adhesion in inhibitory synapse development. Current opinion in neurobiology 18, 77–83 (2008).

26. Y. Iwai, M. Fagiolini, K. Obata, T. K. Hensch, Rapid critical period induction by tonic inhibition in visual cortex. J Neurosci 23, 6695–6702 (2003).

27. F. K. Satterstrom et al., Large-Scale Exome Sequencing Study Implicates Both Developmental and Functional Changes in the Neurobiology of Autism. Cell 180, 568–584 e523 (2020).

28. L. L. Orefice et al., Targeting Peripheral Somatosensory Neurons to Improve Tactile-Related Phenotypes in ASD Models. Cell 178, 867–886 e824 (2019).

29. T. M. DeLorey, P. Sahbaie, E. Hashemi, G. E. Homanics, J. D. Clark, Gabrb3 gene deficient mice exhibit impaired social and exploratory behaviors, deficits in non-selective attention and hypoplasia of cerebellar vermal lobules: a potential model of autism spectrum disorder. Behav Brain Res 187, 207–220 (2008).

30. J. J. LeBlanc, M. Fagiolini, Autism: a "critical period" disorder? Neural Plast 2011, 921680 (2011).

31. J. Michalczyk, A. Milosz, M. Gesek, A. Fornal, Prenatal Diabetes and Obesity: Implications for Autism Spectrum Disorders in Offspring - A Comprehensive Review. Med Sci Monit 30, e945087 (2024).

32. C. M. Vacher, A. Tsompanidis, M. R. Firestein, A. A. Penn, Neuroactive steroid exposure impacts neurodevelopment: Comparison of human and rodent placental contribution. J Neuroendocrinol 10.1111/jne.13489, e13489 (2025).

33. I. H. Greger, L. Khatri, X. Kong, E. B. Ziff, AMPA receptor tetramerization is mediated by Q/R editing. Neuron 40, 763–774 (2003).

34. I. H. Greger, L. Khatri, E. B. Ziff, RNA editing at arg607 controls AMPA receptor exit from the endoplasmic reticulum. Neuron 34, 759–772. (2002).

35. L. Li et al., CASK and FARP localize two classes of post-synaptic ACh receptors thereby promoting cholinergic transmission. PLoS Genet 18, e1010211 (2022).

36. J. S. Packer et al., A lineage-resolved molecular atlas of C. elegans embryogenesis at single-cell resolution. Science 365 (2019).

37. M. E. Boeck et al., The time-resolved transcriptome of C. elegans. Genome Res 26, 1441–1450 (2016).

38. T. Hashimshony, M. Feder, M. Levin, B. K. Hall, I. Yanai, Spatiotemporal transcriptomics reveals the evolutionary history of the endoderm germ layer. Nature 519, 219–222 (2015).

39. G. P. Mullen et al., The Caenorhabditis elegans snf-11 gene encodes a sodium-dependent GABA transporter required for clearance of synaptic GABA. Mol Biol Cell 17, 3021–3030 (2006).

40. G. Jiang et al., A Na+/Cl--coupled GABA transporter, GAT-1, from Caenorhabditis elegans: structural and functional features, specific expression in GABA-ergic neurons, and involvement in muscle function. J Biol Chem 280, 2065–2077 (2005).

41. S. L. McIntire, E. Jorgensen, J. Kaplan, H. R. Horvitz, The GABAergic nervous system of Caenorhabditis elegans. Nature 364, 337–341 (1993).

42. M. Gendrel, E. G. Atlas, O. Hobert, A cellular and regulatory map of the GABAergic nervous system of C. elegans. Elife 5 (2016).

43. J. J. Westmoreland, J. McEwen, B. A. Moore, Y. Jin, B. G. Condie, Conserved function of Caenorhabditis elegans UNC-30 and mouse Pitx2 in controlling GABAergic neuron differentiation. J Neurosci 21, 6810–6819. (2001).

44. O. Hobert, K. Tessmar, G. Ruvkun, The Caenorhabditis elegans lim-6 LIM homeobox gene regulates neurite outgrowth and function of particular GABAergic neurons. Development 126, 1547–1562 (1999).

45. P. Kratsios, A. Stolfi, M. Levine, O. Hobert, Coordinated regulation of cholinergic motor neuron traits through a conserved terminal selector gene. Nat Neurosci 15, 205–214 (2011).

46. C. Rivera et al., The K+/Cl-co-transporter KCC2 renders GABA hyperpolarizing during neuronal maturation. Nature 397, 251–255 (1999).

47. Y. Ben-Ari, Excitatory actions of gaba during development: the nature of the nurture. Nat Rev Neurosci 3, 728–739 (2002).

48. J. E. Tanis, A. Bellemer, J. J. Moresco, B. Forbush, M. R. Koelle, The potassium chloride cotransporter KCC-2 coordinates development of inhibitory neurotransmission and synapse structure in Caenorhabditis elegans. J Neurosci 29, 9943–9954 (2009).

49. A. Bellemer, T. Hirata, M. F. Romero, M. R. Koelle, Two types of chloride transporters are required for GABA(A) receptor-mediated inhibition in C. elegans. EMBO J 30, 1852–1863 (2011).

50. B. Mulcahy et al., Post-embryonic remodeling of the C. elegans motor circuit. Curr Biol 32, 4645–4659 e4643 (2022).

51. S. L. McIntire, R. J. Reimer, K. Schuske, R. H. Edwards, E. M. Jorgensen, Identification and characterization of the vesicular GABA transporter. Nature 389, 870–876 (1997).

52. M. Hammarlund, M. T. Palfreyman, S. Watanabe, S. Olsen, E. M. Jorgensen, Open syntaxin docks synaptic vesicles. PLoS Biol 5, e198 (2007).

53. G. Schiavo et al., Tetanus and botulinum-B neurotoxins block neurotransmitter release by proteolytic cleavage of synaptobrevin. Nature 359, 832–835 (1992).

54. R. M. Weimer et al., Defects in synaptic vesicle docking in unc-18 mutants. Nat Neurosci 6, 1023–1030 (2003).

55. N. X. Tritsch, J. B. Ding, B. L. Sabatini, Dopaminergic neurons inhibit striatal output through non-canonical release of GABA. Nature 490, 262–266 (2012).

56. X. J. Tong, Z. Hu, Y. Liu, D. Anderson, J. M. Kaplan, A network of autism linked genes stabilizes two pools of synaptic GABA(A) receptors. Elife 4, e09648 (2015).

57. X. Zhou et al., The netrin receptor UNC-40/DCC assembles a postsynaptic scaffold and sets the synaptic content of GABAA receptors. Nat Commun 11, 2674 (2020).

58. W. Koh, H. Kwak, E. Cheong, C. J. Lee, GABA tone regulation and its cognitive functions in the brain. Nat Rev Neurosci 24, 523–539 (2023).

59. H. Cheng et al., Phasic/tonic glial GABA differentially transduce for olfactory adaptation and neuronal aging. Neuron 112, 1473–1486 e1476 (2024).

60. B. Graziano et al., Glial KCNQ K(+) channels control neuronal output by regulating GABA release from glia in C. elegans. Neuron 112, 1832–1847 e1837 (2024).

61. V. Riazanski et al., Presynaptic CLC-3 determines quantal size of inhibitory transmission in the hippocampus. Nat Neurosci 14, 487–494 (2011).

62. H. Sun, T. Tsunenari, K. W. Yau, J. Nathans, The vitelliform macular dystrophy protein defines a new family of chloride channels. Proc Natl Acad Sci U S A 99, 4008–4013 (2002).

63. S. Sankaranarayanan, D. De Angelis, J. E. Rothman, T. A. Ryan, The use of pHluorins for optical measurements of presynaptic activity. Biophys J 79, 2199–2208 (2000).

64. G. Miesenbock, D. A. De Angelis, J. E. Rothman, Visualizing secretion and synaptic transmission with pH-sensitive green fluorescent proteins. Nature 394, 192–195 (1998).

65. J. S. Dittman, J. M. Kaplan, Factors regulating the abundance and localization of synaptobrevin in the plasma membrane. Proc Natl Acad Sci U S A 103, 11399–11404 (2006).

66. R. Kohn et al., Expression of multiple UNC-13 proteins in the C. elegans nervous system. Mol. Biol. Cell 11 (2000).

67. M. Nonet, O. Saifee, H. Zhao, J. Rand, L. Wei, Synaptic transmission deficits in Caenorhabditis elegans synaptobrevin mutants. J. Neurosci. 18, 70–80 (1998).

68. A. P. Owji et al., Neurotransmitter-bound bestrophin channel structures reveal small molecule drug targeting sites for disease treatment. Nat Commun 15, 10766 (2024).

69. J. Wang et al., GAD65 tunes the functions of Best1 as a GABA receptor and a neurotransmitter conducting channel. Nat Commun 15, 8051 (2024).

70. M. Di Palma, W. Koh, C. J. Lee, F. Conti, A quantitative analysis of bestrophin 1 cellular localization in mouse cerebral cortex. Acta Physiol (Oxf) 241, e14245 (2025).

71. J. Zhao, L. Gao, S. Nurrish, J. M. Kaplan, Post-synaptic GABA(A) receptors potentiate transmission by recruiting CaV2 channels to their inputs. Cell Rep 42, 113161 (2023).

72. S. G. Brickley, I. Mody, Extrasynaptic GABA(A) receptors: their function in the CNS and implications for disease. Neuron 73, 23–34 (2012).

73. S. Brenner, The genetics of Caenorhabditis elegans. Genetics 77, 71–94 (1974).

74. A. Kumar et al., Dual-view plane illumination microscopy for rapid and spatially isotropic imaging. Nat Protoc 9, 2555–2573 (2014).

75. L. H. Duncan et al., Isotropic Light-Sheet Microscopy and Automated Cell Lineage Analyses to Catalogue Caenorhabditis elegans Embryogenesis with Subcellular Resolution. J Vis Exp 10.3791/59533 (2019).

76. M. E. Boeck et al., Data from “The time-resolved transcriptome of C. elegans.”. https://www.ncbi.nlm.nih.gov/pubmed/27531719.

77. E. L. Ardiel et al., Data from “Stereotyped behavioral maturation and rhythmic quiescence in C. elegans embryos.” https://figshare.com/articles/journal_contribution/2017_04_06_zip/16725349.

