## Supplemental figures and methods for "Immature *C. elegans* motor neurons control early embryo behavior via both synaptic and non-synaptic GABA release"

**This PDF file includes:**

Figures S1 to S6

Table S1

Extended Methods

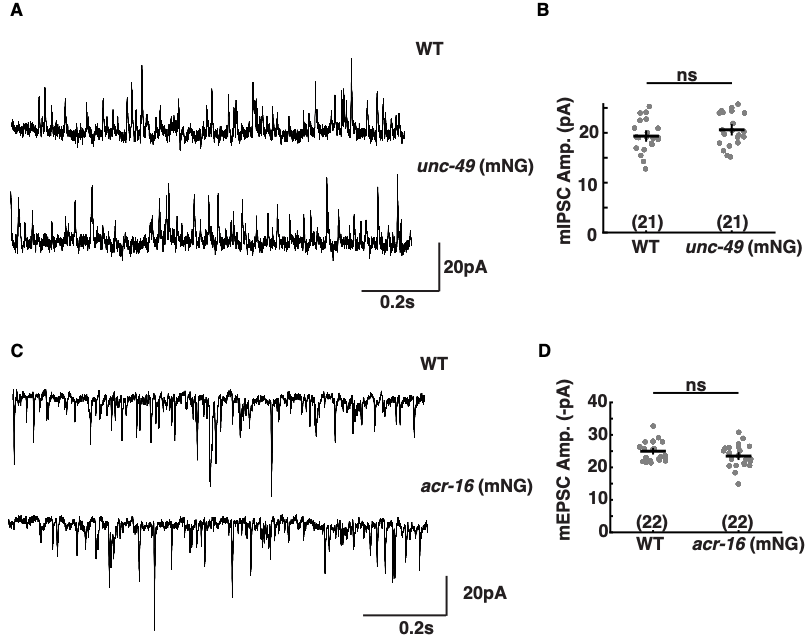

**Figure S1. Supplementary Data for Figure 1.** mNG tags did not significantly alter the function of UNC-49 and ACR-16 channels. Representative traces of adult body muscle miniature inhibitory post-synaptic currents (mIPSCs) (A) and miniature excitatory post-synaptic currents (mEPSCs) (C), mean mIPSC amplitudes (B), and mean mEPSC amplitudes (D) are shown for the indicated genotypes. No significant differences were observed. Error bars indicate SEM. Sample sizes are shown for each genotype.

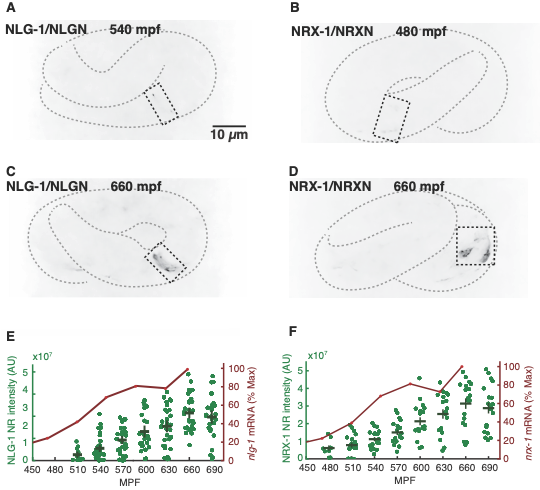

**Figure S2. Supplementary data for Figure 1.** Representative images of early (A,B) and late (C,D) embryos expressing NLG-1/NLGN(nu739 mNG) (A,C) and NRX-1/NRXN (nu742 mNG) are shown. In each case, nerve ring signal (within the dashed rectangles) is observed in the late (C,D) but not early (A,B) embryos. Developmental time point is indicated for each image. (E,F) Mean nerve ring intensity and peak normalized mRNA levels versus developmental time (1, 2) are shown for NLG-1/NLGN (E) and NRX-1/NRXN (F). Error bars indicate SEM.

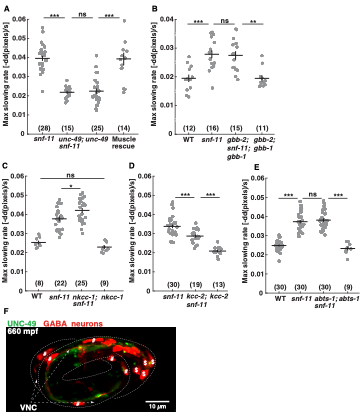

**Figure S3. GABA effects on embryo motion require UNC-49/GABA_A_ receptors.** A) Loss of UNC-49 GABA_A_ receptors eliminates inhibition of flipping in *snf-11* embryos. Inhibition of flipping can be restored via introduction of a transgene driving UNC-49 expression in muscles. B) Loss of metabotropic GABA receptors (*gbb-1; gbb-2* GABA_B_R double mutants) fails to suppress inhibition of flipping in *snf-11* embryos. Motion inhibition in *snf-11* GAT1 embryos is increased in *nkcc-1* mutants (C), is decreased in *kcc-2* mutants (D), and is unaffected in *abts-1* mutants (E). These results suggest that GABA inhibits embryo motion by hyperpolarizing body muscles. Maximal slowing rate is plotted for the indicated genotypes. Sample sizes for each genotype are indicated in each figure panel. Values that differ significantly are indicated (ns, not significant; *, *p* <0.05; ***, *p* <0.001). Error bars indicate SEM. (F) UNC-49(nu829 mNG) fluorescence is observed in ventral and not in dorsal muscles in embryos. The ventral surface of embryos were identified by expressing mChopti in GABA neurons (using the *unc-47* promoter). DD GABA motor neurons (#) run along the ventral side of the body. RME neurons ($) are in the head. Selective UNC-49 expression in ventral muscles explains GABA’s preferential effect on ventral muscle relaxation in *snf-11* GAT1 mutants.

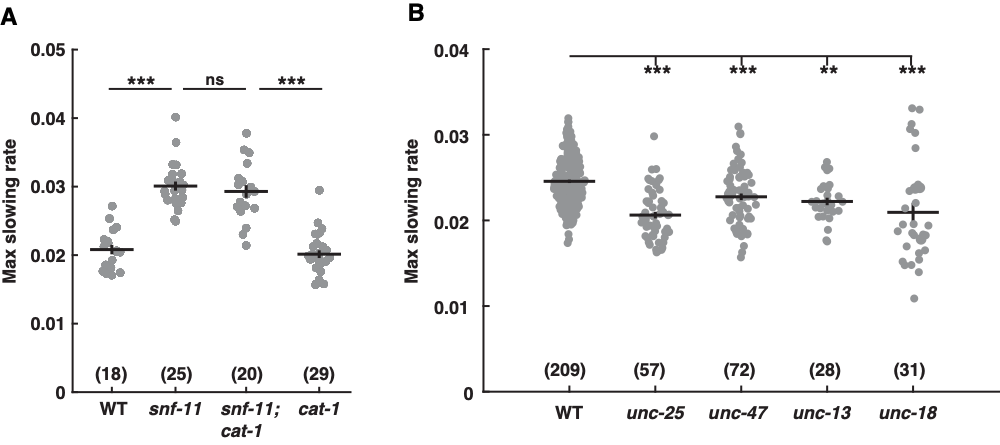

**Figure S4. Supplemental data for Figure 4.** (A) Inactivating CAT-1/VMAT had no effect on embryo motion inhibition in *snf-11* GAT1 mutants. (B) Single mutants lacking UNC-25/GAD, UNC-47/VGAT, UNC-13, and UNC-18 all significantly increased embryo motio. These results indicate that GABA inhibition of embryo motion does not require inactivation of SNF-11/GAT1. Maximal slowing rate of 480-620 mpf embryos is plotted for the indicated genotypes. Sample sizes for each genotype are indicated in each figure panel. Values that differ significantly are indicated (ns, not significant; **, *p* <0.01; ***, *p* <0.001). Error bars indicate SEM.

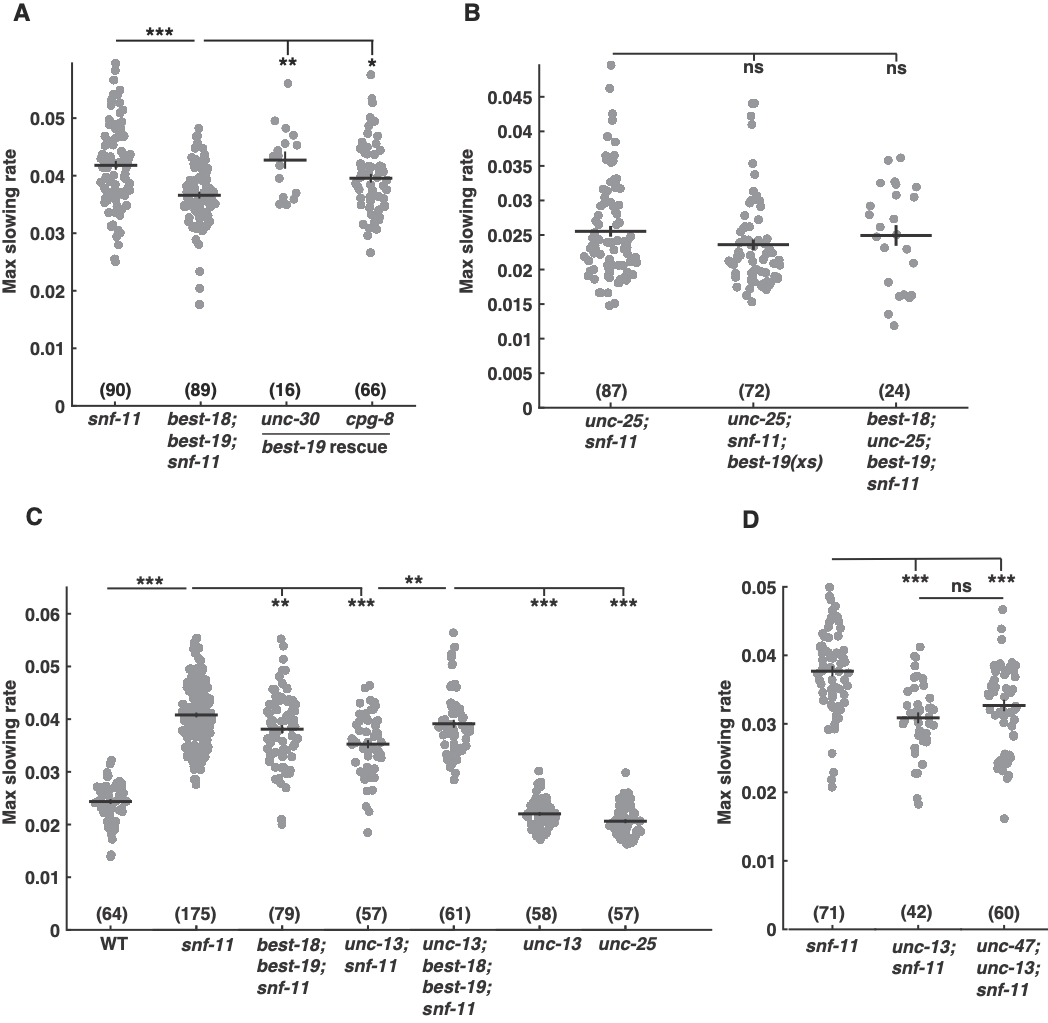

**Figure S5. Bestrophins promote synaptic GABA release from embryonic DD neurons.** (A) Inactivating BEST-18 and BEST-19 significantly decreased motion inhibition in *snf-11* GAT1 mutants and this effect was rescued by transgenes expressing BEST-19 in DD neurons (using the indicated promoters). (B) Bestrophin mutations and over-expression had no effect on motion inhibition of *unc-25; snf-11* double mutants, indicating that bestrophins’ impact on behavior requires GABA signaling. (C) Bestrophin mutations and *unc-13* mutations do not have additive effects on inhibition of embryo motion in *snf-11* GAT1 mutants. Lack of additive defects suggests that Bestrophins promote synaptic GABA transmission. (D) As expected, *unc-13* and *unc-47* VGAT mutations also do not have additive effects on inhibition of embryo motion in *snf-11* mutants, consistent with both acting to promote synaptic GABA release. Maximal slowing rate of 480-620 mpf embryos is plotted for the indicated genotypes. Sample sizes for each genotype are indicated in each figure panel. Values that differ significantly are indicated (ns, not significant; *, *p* <0.05; **, *p* <0.01; ***, *p* <0.001). Error bars indicate SEM.

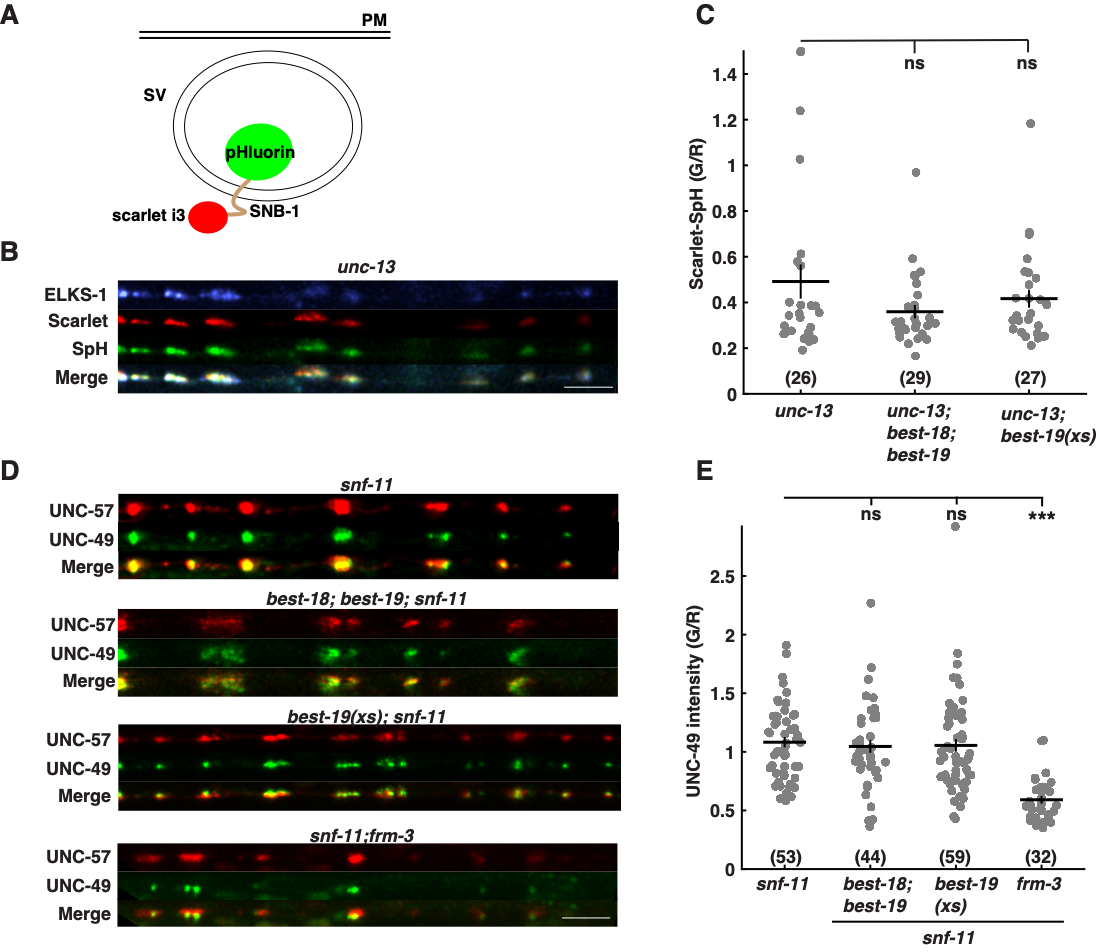

**Figure S6. Supplemental data for Figure 6.** (A-C) Bestrophin mutations and over-expression had no effect on SV acidification. (A) ELKS-1-BFP and Scarlet-SpH were co-expressed in GABA neurons (using the *unc-25* promoter) in *unc-13* mutants, thereby confining Scarlet-SpH to the intracellular pool of SVs. ELKS-1-BFP was used to identify DD neuron active zones in the dorsal nerve cord of adults. Representative images of *unc-13* adults (B) and mean Scarlet-SpH (Green/Red ratio) in the indicated genotypes are shown (C). (D-E) Bestrophin mutations and over-expression had no effect on post-synaptic UNC-49(nu829 mNG) fluorescence at adult DD NMJs. By contrast, *frm-3* FARP mutants significantly decreased post-synaptic UNC-49 levels. DD presynaptic elements were identified by UNC-57-mCherry fluorescence (expressed with the *unc-25* promoter). Representative images (D) and mean post-synaptic UNC-49 intensity (Green/Red ratio) are shown for the indicated genotypes. Values that differ significantly are indicated (ns, not significant; ***, *p* <0.001). Error bars indicate SEM. Scale bars indicate 5 μm.

**Supplementary Table 1. Key Resources Table**

| **Reagent type** | **Source or reference** | **Identifier** |
| --- | --- | --- |
| **Experimental Models:**  **Organisms/Strains** |  |  |
| strain (*C. elegans* ) N2 | CGC | Wormbase ID:N2 |
| strain (*C. elegans* ) KP8564 unc-2(nu569) | Zhao et al. (3) | nu569 contains mNG inserted after codon 2013 of UNC-2B |
| strain (*C. elegans* ) KP11347 unc-49(nu829) | This study | nu829 contains mNG inserted after codon 392 |
| strain (*C. elegans* ) KP10197 acr-16(nu718) | This study | nu718 contains mNG inserted after codon 409 |
| strain (*C. elegans* ) RM2710 snf-11(ok156) | CGC | Wormbase ID:RM2710 |
| strain (*C. elegans* ) KP10834 snf-11(tm625) | NBRP (https://shigen.nig.ac.jp) |  |
| strain (*C. elegans* ) KP10952 snf-11(nu775) wIs51 | This study | wIs51 is SCMp::GFP *+* unc-119(+)  nu775 deletes from 417 bp upstream start codon to the 3’ splice site before exon 5 of *snf-11* |
| strain (*C. elegans* ) KP10953 snf-11(ok156); nuSi620 | This study | nuSi620 is Psbt-1::SNF-11 |
| strain (*C. elegans* ) KP10954 snf-11(ok156); nuSi622 | This study | nuSi622 is Pmyo-3::SNF-11 |
| strain (*C. elegans* ) KP11431 unc-25(nu836) | This study | nu836 deletes from 263 bp upstream of the start codon to 915 bp downstream of the stop codon of *unc-25* |
| strain (*C. elegans* ) KP11432 unc-25(nu836); snf-11(ok156) | This study |  |
| strain (*C. elegans* ) KP10963 ujIs113; snf-11(nu775) wIs51 | This study | wIs51 is SCMp::GFP *+* unc-119(+); ujIs113 is pie-1p::mCherry::H2B::pie-1 3'UTR + nhr-2p::his-24::mCherry::let-858 3'UTR + unc-119(+) |
| strain (*C. elegans* ) LX929 vsIs48 | CGC | Wormbase ID:LX929  vsIs48 is Punc-17::GFP |
| strain (*C. elegans* ) KP11207 vsIs48; snf-11(ok156) | This study | vsIs48 is Punc-17::GFP |
| strain (*C. elegans* ) CB845 unc-30(e191) | CGC | Wormbase ID:CB845 |
| strain (*C. elegans* ) KP10787 unc-30(e191); snf-11(ok156) | This study |  |
| strain (*C. elegans* ) HBR914 lim-6(tm4836) | CGC | Wormbase ID:HBR914 |
| strain (*C. elegans* ) KP10798 snf-11(ok156); lim-6(tm4836) | This study |  |
| strain (*C. elegans* ) KP11257 unc-25(nu808) | This study | nu808 contains mNG inserted after codon 1 |
| strain (*C. elegans* ) KP10913 oxIs12 | Jorgensen Lab | oxIs12 is Punc-47::GFP + lin-15(+) |
| strain (*C. elegans* ) KP6566 gbb-2(tm1165); gbb-1(tm1406) | Dittman and Kaplan (4) |  |
| strain (*C. elegans* ) KP10757 gbb-2(tm1165); snf-11(ok156); gbb-1(tm1406) | This study |  |
| strain (*C. elegans* ) CB407 unc-49(e407) | CGC | Wormbase ID:CB407 |
| strain (*C. elegans* ) KP10759 unc-49(e407); snf-11(ok156) | This study |  |
| strain (*C. elegans* ) KP10973 unc-49(e407); nuSi624; snf-11(ok156) | This study | nuSi624 is Ppat-10::UNC-49B |
| strain (*C. elegans* ) RB1424 nkcc-1(ok156) | CGC | Wormbase ID:RB1424 |
| strain (*C. elegans* ) KP11433 nkcc-1(ok1621); snf-11(ok156) | This study |  |
| strain (*C. elegans* ) LX999 kcc-2(vs132) | CGC | Wormbase ID:LX999 |
| strain (*C. elegans* ) KP10959 kcc-2(vs132); snf-11(nu775) wsIs51 | This study |  |
| strain (*C. elegans* ) RM1381 abts-1(ok1566); snf-11(ok156) | CGC | Wormbase ID:RB1381 |
| strain (*C. elegans* ) KP10877 abts-1(ok1566); snf-11(ok156) | This study |  |
| strain (*C. elegans* ) VC311 unc-47(gk192) | CGC | Wormbase ID:VC311 |
| strain (*C. elegans* ) KP10828 unc-47(gk192); snf-11(ok156) | This study |  |
| strain (*C. elegans* ) CB1111 cat-1(e1111) | CGC | Wormbase ID:CB1111 |
| strain (*C. elegans* ) KP10958 cat-1(e1111); snf-11(nu775) wsIs51 | This study |  |
| strain (*C. elegans* ) BC168 unc-13(s69) | CGC | Wormbase ID:BC168 |
| strain (*C. elegans* ) KP10799 unc-13(s69); snf-11(ok156) | This study |  |
| strain (*C. elegans* ) CB81 unc-18(e81) | CGC | Wormbase ID:CB81 |
| strain (*C. elegans* ) KP10878 snf-11(ok156); unc-18(e81) | This study |  |
| strain (*C. elegans* ) KP10940 syIs396 syIs420; snf-11(ok156) | This study | syIs396 is Punc-47::NLS::NLS::GAL4SK::VP64::let-858 3'UTR + unc-122p::RFP + 1kb DNA ladder (NEB) + Pttx-3::RFP + 1kb DNA ladder(NEB)  syIs420 is 15xUAS::*pes-10*::TETX::*let-858* 3'UTR + P*myo-2*::NLS::GFP + pBlueScript |
| strain (*C. elegans* ) VC1288 frm-3(gk585) | CGC | Wormbase ID:VC1288 |
| strain (*C. elegans* ) KP11434 frm-3(gk585); snf-11(ok156) | This study |  |
| strain (*C. elegans* ) KP111349 best-18(gk5761); snf-11(ok156) | This study |  |
| strain (*C. elegans* ) KP11360 best-19(nu835); snf-11(ok156) | This study |  |
| strain (*C. elegans* ) KP11363 best-18(gk5761); best-19(nu835); snf-11(ok156) | This study |  |
| strain (*C. elegans* ) KP11436 snf-11(ok156); nuSi730 | This study | nuSi730 is Punc-25::BEST-19 |
| Strain (*C. elegans* ) KP11435  best-18(gk5761) unc-47(nu855); best-19(nu835); snf-11(ok156) | This study |  |
| Strain (*C. elegans* ) KP11440 unc-47(nu855); snf-11(ok156) | This study |  |
| Strain (*C. elegans* ) KP11464 best-18(gk5761) unc-25(nu836); best-19(nu835); snf-11(ok156) | This study |  |
| Strain (*C. elegans*) KP11462 unc-25(nu836); snf-11(ok156); nuSi730 | This study | nuSi730 is Punc-25::BEST-19 |
| Strain (*C. elegans*) KP11466 unc-40(e271); snf-11(nu775) wIs51 | This study | wIs51 is SCMp::GFP *+* unc-119(+); |
| Strain (*C. elegans*) CB271 unc-40(e271) | CGC | Wormbase ID:CB271 |
| Strain (*C. elegans*) KP11494 unc-3(e151); snf-11(ok156) | This study |  |
| Strain (*C. elegans*) CB151 unc-3(e151) | CGC | Wormbase ID:CB151 |
| Strain (*C. elegans*) KP11359 unc-25(nu808); snf-11(ok156) | This study | nu808 contains mNG inserted after codon 1 |
| Strain (*C. elegans*) KP11495 oxIs12; snf-11(ok156) | This study | oxIs12 is Punc-47::GFP + lin-15(+) |
| Strain (*C. elegans*) KP11437 unc-13(s69); best-18(gk5761); best-19(nu835); snf-11(ok156) | This study |  |
| Strain (*C. elegans*) KP11497 unc-13(s69); unc-47(nu855); snf-11(ok156) | This study |  |
| Strain (*C. elegans)* KP11498 unc-13(s69); nuSi737 | This study | nuSi737 is Punc-25::scarlet i3:snb-1:pHluorin SL2 BFP:elks-1 |
| Strain (*C. elegans)* KP11499 unc-13(s69); nuSi730; nuSi737 | This study |  |
| Strain (*C. elegans)* KP11500 unc-13(s69); best-18(nu863); best-19(nu835) | This study | best-19(nu863) is best-18(gk5761) exposed to germline-expressed CRE to excise the loxP Pmyo-2::GFP::unc-54 3’ UTR + Prps-27::neoR::unc-54 3’ UTR + loxP cassette |
| Strain (*C. elegans)* KP11501 nuSi285; unc-49(nu829); snf-11(ok156) | This study | nuSi285 is Punc-47:GFP1-10:SL2:unc-57:mCherry:SL2:mTaqBFP2 |
| Strain (*C. elegans)* KP11502 nuSi285; best-18(nu863) unc-49(nu829); best-19(nu835); snf-11(ok156) | This study |  |
| Strain (*C. elegans)* KP11503 nuSi285; nuSi730 unc-49(nu829); snf-11(ok156) | This study |  |
| Strain (*C. elegans)* KP11500 frm-3(gk585); nuSi285; unc-49(nu829); snf-11(ok156) | This study |  |
| Strain (*C. elegans*) KP10492 nlg-1(nu739) | Zhao et al. (3) | nu739 contains mNeonGreen inserted at codon 791 of NLG-1A |
| Strain (*C. elegans*) KP10506 nrx-1(nu742) | This study | nu742 contains mNeonGreen inserted at codon 1515 of NRX-1A |
| Strain (*C. elegans*) KP11510 nuSi742; best-18(gk5761); best-19(nu835); snf-11(ok156) | This study | nuSi742 is Punc-30:BEST-19 |
| Strain (*C. elegans*) KP11516 nuSi748; best-18(gk5761); best-19(nu835); snf-11(ok156) | This study | nuSi748 is Pcpg-8:BEST-19 |
| Strain (*C. elegans)* KP11535 otIs374; unc-49(nu829) | This study | otIs374 is Punc-47::mChopti::unc-54 3’ UTR + pha-1(+) |
| **Oligonucleotides: Guide RNAs** |  |  |
| aactacagacgtctcacaac AGG | IDT | guide RNA for acr-16 neonGreen insertion |
| ttttctgggatctccatcaa TGG | IDT | guide RNA for unc-49 neonGreen insertion |
| aataagggacactgtgctag TGG | IDT | guide RNA for snf-11 deletion allele (nu775), 5’ end |
| taccgggtagtcataattgt AGG | IDT | guide RNA for snf-11 deletion allele (nu775), 3’ end |
| gtttcgtgacgagtcttcaa AGG | IDT | guide RNA for unc-25 loxP insertion (nu802), 5’ end |
| tagtagctttgaccactttc AGG | IDT | guide RNA for unc-25 loxP insertion (nu802), 3’ end |
| aactgagcttttccctattc CGG | IDT | guide RNA for unc-25 neonGreen insertion |
| cacgccatctcaactgtaat CGG | IDT | guide RNA for best-19 deletion allele (nu835), 5’ end |
| tggtgatgacgtggaaaaga AGG | IDT | guide RNA for best-19 deletion allele (nu835), 3’ end |
| ttcgctggactactggaatc AGG | IDT | guide RNA for unc-47 deletion allele (nu855), 5’ end |
| tggtgtctacttctcatcaa TGG | IDT | guide RNA for unc-47 deletion allele (nu855), 3’ end |
| acctgtatctcttccaatgt CGG | IDT | guide RNA for nlg-1 neonGreen insertion allele (nu739) |
| **Recombinant DNA** |  |  |
| Plasmid: KP#4532 Ppat-10 UNC-49B miniMOS hyg | This study | Expresses UNC-49B in body wall muscles |
| Plasmid: KP#4600 Pmyo-3 SNF-11 miniMOS hyg | This study | Expresses SNF-11 in body wall muscles |
| Plasmid: KP#4601 Psbt-1 SNF-11 miniMOS hyg | This study | Expresses SNF-11 in neurons |
| Plasmid: KP#4602 Punc-25 BEST-19 miniMOS hyg | This study | Expresses BEST-19 in GABA neurons |
| Plasmid: KP#4603 Punc-30 BEST-19 miniMOS hyg | This study | Expresses BEST-19 in GABA motor neurons |
| Plasmid: KP#4604 Pcpg-8 BEST-19 miniMOS hyg | This study | Expresses BEST-19 in a subset of GABA neurons |
| Plasmid: KP#4605 Punc-25 scarlet i3 snb-1 pHluorin SL2 BFP elks-1 | This study | Expresses Scarlet i3:SynaptopHluorin and BFP:ELKS-1 in GABA neurons |
| **Chemicals/Drugs** |  |  |
| Poly-L-lysine | Sigma-Aldrich | Catalog #: P2636 |
| Polybead microspheres 0.10 μm | Polysciences | Catalog #: 00876-15 |
| **Software and Algorithms** |  |  |
| NIS Elements | Nikon | https://www.microscope.healthcare.nikon.com/ |
| ImageJ/FIJI | NIH | https://fiji.sc |
| Micro-Manager | Edelstein et al. (5) | https://micro-manager.org |
| Matlab R2024b | Mathworks | https://www.mathworks.com/products/matlab.html |
| pClamp 10 | Molecular Devices | https://www.moleculardevices.com |

**Extended Methods.**

**Transgenes.** All plasmids were built from pCFJ910 (6). For *snf-11* rescue experiments, a *snf-11* cDNA was expressed in muscle with a 2600 bp *myo-3* promoter (plasmid KP#4600) or in neurons with a 951 bp *sbt-1* promoter (plasmid KP#4601). For *unc-49* rescue, an *unc-49B* cDNA was expressed in muscle with a *pat-10* promoter (plasmid KP#4532). For *best-19* over-expression in GABA neurons, a *best-19* cDNA was expressed with an *unc-25* promoter (plasmid KP#4602).

The miniMOS method(6) was used to make the following single copy transgenes: *nuSi624[Ppat-10:unc-49B], nuSi622[Pmyo-3:snf-11]*, *nuSi620*[*Psbt-1:snf-11*], *nuSi730*[*Punc-25:best-19*], *nuSi742[Punc-30:best-19]*, *nuSi748[Pcpg-8:best-19]*, and *nuSi737[Punc-25:scarlet i3:snb-1:pHluorin:SL2:BFP:elks-1]*.

**CRISPR alleles.** CRISPR alleles were isolated as described (7). Briefly, guide RNAs and Cas9 protein, as well as single stranded ULTRAMER oligonucleotide repair templates shorter than 200 bp, were obtained from IDT. The pRF4 *rol-6*(gf) was included in the injection mix. Injected animals were singled and ~100 F1 progeny were singled from plates containing rollers. Progeny of isolated F1’s were screened by PCR for the expected genome edits. Sequences of repair oligonucleotides and of genome edits are provided for each CRISPR allele in the Key Resources Table.

CRISPR was used to generate a floxed *unc-25(nu802*) allele in which LoxP sites flanking the coding sequence were inserted into the *unc-25* locus. The *unc-25(nu836*) null allele was isolated by expressing the CRE recombinase in the germline of *unc-25*(*nu802* FLOX) mutants. Injected animals were singled and F1 progeny were screened for the expected change. The *unc-25(nu836*) allele deletes the entire *unc-25* coding sequence.

The *best-19*(*nu835*) CRISPR allele fully deletes exons 2-8 and portions of exons 1 and 9. The *snf-11*(*nu775*) CRISPR allele deletes all of exons 1-4 and the majority of exon 5. The *unc-47(nu855)* CRISPR allele fully deletes exons 2-6 and portions of exons 1 and 7.

To fluorescently tag *unc-25*, CRISPR was used to insert mNG at the amino-terminus of the endogenous *unc-25* locus. To fluorescently tag *acr-16*, CRISPR was used to insert mNG into the ninth exon of the endogenous *acr-16* locus. To fluorescently tag *unc-49*, CRISPR was used to insert mNG into the tenth exon of *unc-49B.* To fluorescently tag *nlg-1*, CRISPR was used to insert mNG into the last exon of the *nlg-1A.*

**Brightfield motion assay.**

Assays were performed as described in Ardiel et al. (2022) (8). Briefly, gravid day 1 adults were dissected in an M9 filled glass bottom dish (MatTek Corp., P35G-1.5–20 C) and embryos were arrayed on poly-L-lysine (0.1 mg/mL). Embryos were imaged on an inverted microscope (Zeiss, Axiovert 100) with a 5x, 0.25 NA objective. Using a CCD camera (CoolSNAP HQ2) 16 bit images with pixel size of 1.29 µm were acquired with an exposure time of 10ms at 1 Hz using Micro-Manager control software. Transmitted light was calibrated to produce background pixel intensities of approximately 1,500 AU. All images were acquired at room temperature.

To quantify embryo motion, pixel intensities were collected from a 21x21 pixel box within each embryo. Frame-to-frame pixel intensity changes larger than a 100 AU threshold were counted. Onset of twitching and hatching were detected, and frames in which new hatchlings swam into the field of view were excluded, as previously described (8). Embryonic age was determined by measuring time following twitch onset, which was defined as 430 mpf. Minutes post fertilization values for developmental landmarks (such as hatch time) exhibit some day-to-day variation, most likely due to subtle differences in ambient temperature.

**diSpim imaging and posture analysis**

diSpim imaging was performed as described in Ardiel et al. (2022) (8) from ~90 to 180 minutes after twitch onset, as estimated based on observation of embryos under a dissecting microscope. Embryos were dissected as described above. On a diSPIM,(9) a pair of perpendicular water-dipping, long-working distance objectives (40 x, 0.8 NA) were used for brightfield and fluorescence imaging. Acquisition was controlled using Micro-Manager’s diSPIM plugin (<https://micro-manager.org/>). Imaging was done in iSPIM mode, using a single view. A two-dimensional MEMS mirror in the diSPIM scanhead swept a 488 nm laser beam to generate a light sheet and defined image volume. The camera (pco.edge 4.2) chip readout direction was oriented along the embryo’s minor axis to maximize speed. The imaging objective translated in step with the light sheet to produce 36 planes 1.2 µm apart, with exposure times of 4 ms per plane and volumes acquired at 3 Hz. 4X binning to reduce file sizes produced 16 bit images with 0.65 µm pixel sizes. Detailed protocols for diSPIM embryo imaging can be found in Duncan et al. (2019) (10).

*Snf-11* postures were built using *wIs51[SCM*p::GFP + *unc-119*(+)], which expresses GFP in hypodermal seam cell nuclei. Seam cell fluorescence produced by the *wIs51* transgene was too dim to accurately track postures before ~570 mpf. A large kernel 3D U-Net trained on an average of the Dice coefficient and binary cross-entropy loss functions was used to segment images, as described in detail in Ardiel et al. (2022).(8) Seam cells were manually positioned along the body axis by observation of several frames to produce a seed volume. From there, seam cell tracking was performed using a Global Nearest Neighbor (GNN) algorithm as implemented in Matlab. Cell identifications were manually corrected when tracking errors became apparent. Dorsoventral bends were computed from the midpoint along the spline connecting neighboring seam cells and the corresponding midpoint on the opposite side of the body. See Ardiel et al. (2022) (8) for details.

**Fluorescence imaging**

Embryos were dissected from gravid day 1 adults as described under “Brightfield Motion Assay” and synchronized at the 2-cell stage. Embryos were then moved to an inverted confocal microscope (Nikon, Eclipse Ti2) and imaged using a 60X/1.49NA objective with the resonance scanner. Volumes were acquired using resonant scanning and a piezo stage stepping 150 nm between 269 image planes. This generated images of 512x256 pixels (0.14 µm pixels) in the 488nm laser channel. All images were acquired at room temperature. Image denoising was performed using the Denoise.AI algorithm (Nikon). All subsequent image analyses were done using FIJI.

**Nerve ring ion channel imaging and quantitation**

Following synchronization, embryos were imaged as described under “Brightfield Motion Assay” until first twitch. Volumes were acquired using resonant scanning and a piezo stage stepping 150 nm between 269 image planes. This generated images of 512x256 pixels (0.14 µm pixels) in the 488nm laser channel. One volume of each embryo was taken every thirty minutes for 6 hours. The volume of nerve ring fluorescent signals was quantified as follows: images were thresholded at 40 AU, the nerve ring region of interest was identified by manual inspection, and the volume of thresholded nerve ring fluorescence was measured using the FIJI 3D Objects Counter function. Embryos lacking nerve ring signal were scored as 0 area. Embryos that moved during acquisition were excluded from the analysis.

**Dorsoventral bend bias assay**

Following synchronization, embryos were imaged as described under “Brightfield Motion Assay” until first twitch. Volumes were acquired using resonant scanning and a piezo stage stepping 150 nm between 215 image planes. This generated images of 512x256 pixels (0.14 µm pixels) in the 488nm laser channel. One volume of each embryo was taken every five minutes for 30 minutes. Embryos between 540-600 mpf were identified by time of first twitch. Dorsal or ventral bend was manually assessed based on the vsIs48 [*Punc-17:GFP*] marker, which labels cholinergic neurons in the head and along the ventral cord. Differences between strains were assessed using the Chi-square test.

**DD neurite morphology assay**

Following synchronization, embryos were imaged as described under “Brightfield Motion Assay” until first twitch. Volumes were acquired using resonant scanning and a piezo stage stepping 150 nm between 269 image planes. This generated images of 512x256 pixels (0.14 µm pixels) in the 488nm laser channel. One volume of each embryo was taken every 15 minutes for 2.5 hours. DD neurite outgrowth was manually assessed.

**DD UNC-25:mNG localization assay**

Following synchronization, embryos were imaged as described under “Brightfield Motion Assay” until first twitch. Volumes were acquired using resonant scanning and a piezo stage stepping 150 nm between 269 image planes. This generated images of 512x256 pixels (0.14 µm pixels) in the 488nm laser channel. One volume of each embryo was taken every 15 minutes for 2.5 hours. Cell identification and subcellular localization were manually assessed.

### **Muscle mIPSC recordings**

Whole-cell patch-clamp measurements were performed using an Axopatch 200B amplifier with pClamp 10 software (Molecular Devices). The data were sampled at 10 kHz and filtered at 5 kHz. Body muscle IPSCs were recorded as previously described (Tong et al., 2015). Dissected adults were superfused in an extracellular solution containing 127 mM NaCl, 5 mM KCl, 26 mM NaHCO_3_, 1.25 mM NaH_2_PO_4_, 10 mM glucose, 5mM sucrose, 1 mM CaCl_2_, and 4 mM MgCl_2_, bubbled with 5% CO_2_, 95% O_2_ at 22°C. The pipette solution contained 105 mM CH_3_O_3_SCs, 10 mM CsCl, 15 mM CsF, 4mM MgCl_2_, 5mM EGTA, 0.25mM CaCl_2_, 10mM HEPES, and 5mM Na_2_ATP, 1mM Na_2_GTP adjusted to pH 7.2 using CsOH. Whole-cell recordings were carried out at 0mV to record mIPSCs.

**Fluorescence imaging of adult worms**

Day 1 adult worms were immobilized on 10% agarose pads with 3 uL of 0.1 μm diameter polystyrene microspheres (Polysciences 00876-15, 2.5% w/v suspension). The dorsal nerve cord just anterior to the vulva was imaged. Images were taken with a Nikon A1R confocal using a 60X/1.49 NA oil objective. Image volumes spanning the dorsal nerve cord were collected (60-90 planes/volume, 0.15 um between planes, and 0.1 mm/pixel). Maximum intensity projections for each volume were auto-thresholded and puncta were identified as round fluorescent objects (area >0.1 μm^2^) using analysis of particles. Mean fluorescent intensity in each punctum was analyzed in the raw images. All image analysis was done using FIJI.

**UNC-49 puncta analysis**

Pre-synaptic regions of interest (ROIs) were identified by localization of an mCherry-tagged synaptic vesicle marker (UNC-57 Endophilin) expressed in the GABA neurons. Using a CRISPR-tagged UNC-49 allele, the intensity of endogenously tagged UNC-49 in the UNC-57 ROIs was quantified.

**Synaptic vesicle pH quantification**

Synaptic vesicle pH was assessed using a transgenic operon including scarlet i3- and pHluorin-tagged *snb-1* Synaptobrevin and BFP-tagged *elks-1* ERC expressed in the GABA neurons. Pre-synaptic regions of interest (ROIs) were identified by localization of BFP:ELKS-1. The intensity of pHluorin and Scarlet i3 fluorescence in the BFP:ELKS-1 ROI was quantified.

**Quantitation and Statistical Analysis**

For normally distributed data, significant differences were assessed with unpaired t tests (for 2 groups) or one way ANOVA with post-hoc Tukey-Kramer multiple comparisons test (for >2 groups). For non-normal data, differences were assessed by Mann-Whitney (2 groups) or Kruskal-Wallis test with post-hoc Dunn’s multiple comparisons test (>2 groups). For categorical data, differences were assessed by Ch-squared test, with Benjamini-Hochberg procedure applied for false discovery rate where applicable. For cumulative probability distributions, pairwise Kolmogorov-Smirnov tests were used to assess differences. Data graphing and statistics were performed in Matlab 2024b. No statistical method was used to select sample sizes. For brightfield imaging studies, data points represent fastest reduction in movement between 480-620 mpf (methodology described above) in individual embryos (which were considered biological replicates). Details for each experiment can be found in figure legends. For quantitative fluorescence imaging studies, data points represent volume above an intensity threshold in individual embryos (which were considered biological replicates). Embryos that did not hatch during image acquisition, or that were knocked out of position, were excluded from brightfield motion assay analysis. Embryos that moved during volume acquisition were excluded from fluorescence imaging analysis.

**References**

1.	M. E. Boeck *et al.*, The time-resolved transcriptome of C. elegans. *Genome Res* **26**, 1441-1450 (2016).

2.	T. Hashimshony, M. Feder, M. Levin, B. K. Hall, I. Yanai, Spatiotemporal transcriptomics reveals the evolutionary history of the endoderm germ layer. *Nature* **519**, 219-222 (2015).

3.	J. Zhao, L. Gao, S. Nurrish, J. M. Kaplan, Post-synaptic GABA(A) receptors potentiate transmission by recruiting CaV2 channels to their inputs. *Cell Rep* **42**, 113161 (2023).

4.	J. S. Dittman, J. Kaplan, Behavioral impact of neurotransmitter-gated GPCRs: Muscarinic and GABAB receptors regulate C. elegans locomotion. *Journal of Neuroscience* **28**, 7104-7112 (2008).

5.	A. D. Edelstein *et al.*, Advanced methods of microscope control using muManager software. *J Biol Methods* **1** (2014).

6.	C. Frokjaer-Jensen *et al.*, Random and targeted transgene insertion in Caenorhabditis elegans using a modified Mos1 transposon. *Nat Methods* **11**, 529-534 (2014).

7.	K. S. Ghanta, C. C. Mello, Melting dsDNA Donor Molecules Greatly Improves Precision Genome Editing in Caenorhabditis elegans. *Genetics* **216**, 643-650 (2020).

8.	E. L. Ardiel *et al.*, Stereotyped behavioral maturation and rhythmic quiescence in C. elegans embryos. *Elife* **11** (2022).

9.	A. Kumar *et al.*, Dual-view plane illumination microscopy for rapid and spatially isotropic imaging. *Nat Protoc* **9**, 2555-2573 (2014).

10.	L. H. Duncan *et al.*, Isotropic Light-Sheet Microscopy and Automated Cell Lineage Analyses to Catalogue Caenorhabditis elegans Embryogenesis with Subcellular Resolution. *J Vis Exp* 10.3791/59533 (2019).
